# The Role of Hypermutation and Collateral Sensitivity in Antimicrobial Resistance Diversity of *Pseudomonas aeruginosa* Populations in Cystic Fibrosis Lung Infection

**DOI:** 10.1101/2023.06.14.544983

**Authors:** Jelly Vanderwoude, Sheyda Azimi, Timothy D. Read, Stephen P. Diggle

**Affiliations:** Center for Microbial Dynamics and Infection, School of Biological Sciences, Georgia Institute of Technology, Atlanta, GA, USA; School of Biology, Georgia State University, Atlanta, GA, USA; Division of Infectious Diseases, Department of Medicine, Emory University School of Medicine, Atlanta, GA, USA

**Keywords:** cystic fibrosis, population heterogeneity, antibiotic resistance, hypermutation, evolution

## Abstract

*Pseudomonas aeruginosa* is an opportunistic pathogen which causes chronic, drug-resistant lung infections in cystic fibrosis (CF) patients. In this study, we explore the role of genomic diversification and evolutionary trade-offs in antimicrobial resistance (AMR) diversity within *P. aeruginosa* populations sourced from CF lung infections. We analyzed 300 clinical isolates from four CF patients (75 per patient), and found that genomic diversity is not a consistent indicator of phenotypic AMR diversity. Remarkably, some genetically less diverse populations showed AMR diversity comparable to those with significantly more genetic variation. We also observed that hypermutator strains frequently exhibited increased sensitivity to antimicrobials, contradicting expectations from their treatment histories. Investigating potential evolutionary trade-offs, we found no substantial evidence of collateral sensitivity among aminoglycoside, beta-lactam, or fluoroquinolone antibiotics, nor did we observe trade-offs between AMR and growth in conditions mimicking CF sputum. Our findings suggest that (i) genomic diversity is not a prerequisite for phenotypic AMR diversity; (ii) hypermutator populations may develop increased antimicrobial sensitivity under selection pressure; (iii) collateral sensitivity is not a prominent feature in CF strains, and (iv) resistance to a single antibiotic does not necessarily lead to significant fitness costs. These insights challenge prevailing assumptions about AMR evolution in chronic infections, emphasizing the complexity of bacterial adaptation during infection.

**Importance:** Upon infection in the cystic fibrosis (CF) lung, *Pseudomonas aeruginosa* rapidly acquires genetic mutations, especially in genes involved in antimicrobial resistance (AMR), often resulting in diverse, treatment-resistant populations. However, the role of bacterial population diversity within the context of chronic infection is still poorly understood. In this study, we found that hypermutator strains of *P. aeruginosa* in the CF lung undergoing treatment with tobramycin evolved increased sensitivity to tobramycin relative to non-hypermutators within the same population. This finding suggests that antimicrobial treatment may only exert weak selection pressure on *P. aeruginosa* populations in the CF lung. We further found no evidence for collateral sensitivity in these clinical populations, suggesting that collateral sensitivity may not be a robust, naturally occurring phenomenon for this microbe.

**Preprint servers:** This manuscript has been submitted as a preprint to bioRxiv

## Introduction

*Pseudomonas aeruginosa* is a dominant bacterial pathogen in chronic infections of the airways of adults with cystic fibrosis (CF), a genetic disorder that results in thickened mucus, persistent lung infection, and progressive decline in lung function [1, 2]. *P. aeruginosa* has multiple intrinsic and acquired mechanisms of antimicrobial resistance (AMR), with clinical strains sometimes displaying multi-drug resistance (MDR). While antibiotic treatment can be effective against early-stage, transient *P. aeruginosa* infections, in the case of chronic infections, antibiotic regimens ameliorate patient symptoms and prolong life but ultimately fail to eradicate *P. aeruginosa* from the CF lung [3]. This is largely due to the microaerophilic environment of the CF lung leading to slow growth and the viscous mucosal matrix hindering drug penetration [4, 5]. Treatment failure may additionally result from the high degree of phenotypic and genomic heterogeneity that naturally evolves in *P. aeruginosa* populations inhabiting CF airways [6], allowing the population to exploit various pathways of resistance and for the emergence of rare clones that evade treatment and re-establish infection afterwards [7, 8]. Most individuals with CF are initially infected by a single environmental or transmissible epidemic strain of *P. aeruginosa*, which then diversifies in the CF lung over the course of many years of infection [9]. Mutations in DNA mismatch repair (MMR) mechanisms act as a catalyst for this diversification, potentially providing an evolutionary advantage in an environment that demands rapid adaptation for survival, though potentially at a fitness cost [10, 11].

Maintaining diversity in populations can be advantageous for bet-hedging in a complex infection environment where there are a multitude of external stressors such as competing microbiota, antibiotic exposure, and host immune responses. Heterogeneity in populations may develop as individual members of the population evolve specialized functions to occupy different ecological niches [12], however, adaptations to a particular niche may come at an expense to other energetically costly traits (i.e., fitness costs) [13, 14]. The vast diversity of *P. aeruginosa* in CF lung infection suggests that individual isolates within the population could have different specializations resulting in trade-offs with other traits. Of particular interest to researchers is collateral sensitivity— increased sensitivity to one antimicrobial as a trade-off with increased resistance to another— as a potential avenue for targeting drug-resistant populations using combination therapy or antibiotic cycling. Although collateral sensitivity has been evolved *in vitro* [15–19], it remains to be determined whether collateral sensitivity is robust across naturally occurring clinical populations of *P. aeruginosa*.

Despite *P. aeruginosa* population diversity in the CF lung being widely accepted, this diversity is often overlooked. Within-host adaptations of *P. aeruginosa* to the CF lung have previously been investigated and described, primarily via longitudinal single-isolate sampling [20–30]. Longitudinal sampling of single or small subsets of isolates from a population only reflects a fraction of the total evolutionary pathways exhibited within a population and may result in significant underestimation of the diversity of antimicrobial susceptibility profiles. As population diversity may impact infection outcomes via heteroresistance [31], microbial social interactions [32, 33], or the ability of a population to survive evolutionary bottlenecks [3], this warrants a shift in our sampling and susceptibility testing of chronic microbial infections to reflect our understanding of them as complex, dynamic populations. A few studies have thoroughly investigated population diversity in this infection context, in which their analyses were focused on (i) phenotypic diversity [34–38]; (ii) genetic analyses via pooled population sequencing [39, 40]; or (iii) both extensive sequencing and phenotyping, but lacking analysis linking the two at the isolate-level [6]. As a result, we still have an incomplete understanding of how genomic diversification drives AMR heterogeneity within a population, and what trade-offs are involved in these evolutionary processes.

Here, we investigated genomic and AMR diversity for chronic *P. aeruginosa* lung populations in four unique individuals with CF. We first sought to test whether genomic diversity is a strong predictor of phenotypic diversity in AMR within a population. With the rapid advances in sequencing technology, researchers are already investigating methods to replace time-consuming antimicrobial susceptibility testing (AST) with sequencing as a diagnostic tool [41]. As such, our goal was to determine the viability of predicting AMR phenotypic diversity from genomic population diversity in a manner that could easily be translated to the clinic. We further explored the role that hypermutation plays in driving resistance, specific links between genotype and phenotype at the isolate-level, and enrichments in mutations and gene content changes relevant to AMR. Lastly, we searched for evidence that resistance to one antimicrobial may trade-off with sensitivity to other antimicrobials and fitness in a CF-like environment.

## Methods

### Cohort selection and strain isolation

We selected four adult individuals, aged 24-31 years, for this study from a cohort of CF patients at Emory University in Atlanta who had been chronically infected with *P. aeruginosa* for 10-15 years at the time of sampling. From each patient, we collected and processed a single expectorated sputum sample. We processed sputum by supplementing each sample with 5 ml synthetic cystic fibrosis medium (SCFM) [42] and autoclaved glass beads, homogenizing the mixture via vortexing for 2 mins, centrifuging the homogenized sputum mixture for 4 mins at ∼3,300 x *g*, removing the supernatant, and conducting a 10x serial dilution of cell pellet re-suspended in phosphate buffered saline to streak on *Pseudomonas* isolation agar (PIA) plates. These plates were incubated at 37℃ overnight, then at room temperature for up to 72 h. From each expectorated sputum sample, we randomly isolated 75 *P. aeruginosa* colonies for a total of 300 isolates. These isolates were confirmed to be *P. aeruginosa* using 16S rRNA gene amplification before proceeding with whole genome sequencing.

### Whole genome sequencing

To conduct sequencing, we first grew all 300 isolates overnight in 15 ml conical tubes in lysogeny broth (LB) at 37℃ with shaking at 200 rpm. We extracted DNA from these cultures using the Promega Wizard Genomic DNA Purification Kit according to the manufacturer’s instructions. We prepared sequencing libraries using the Nextera XT DNA Library Preparation Kit and used the Illumina Novaseq platform to obtain 250 bp paired-end reads for a mean coverage of 70x. 28 samples either failed or did not meet the minimum sequencing coverage or quality requirements, so we re-sequenced these using the Illumina MiSeq platform for 250 bp paired-end reads and combined the reads from both sequencing runs to analyze these 28 samples. We randomly selected one isolate from each patient to serve as the reference strain for the other 74 isolates isolated from that patient. For these reference isolates, we additionally obtained Oxford Nanopore long read sequences through the Microbial Genome Sequencing Center (GridION Flow Cell chemistry type R9.4.1 with Guppy high accuracy base calling v4.2.2) at 35x coverage.

### Multi-locus sequence typing

Our multi-locus sequence typing was implemented in Bactopia v1.6.5 [43], which employs the PubMLST.org schema [44].

### Constructing annotated reference assemblies

We used Unicyler v0.5.0 [45] to create long-read assemblies for the four reference isolates. We then conducted one round of long-read polishing on these assemblies using Medaka v1.0.3 [46], which produced preliminary consensus sequences. We conducted quality control on all 300 Illumina reads using the Bactopia v1.6.5 [43] pipeline. We conducted two further short-read assembly polishing steps on the long-read assemblies by aligning the quality-adjusted short reads of each of the four reference isolates to its respective consensus sequence using Polypolish v0.5.0 [47] and Pilon v1.24 [48]. We validated the final consensus sequences by mapping the Illumina reads of each reference to its respective assembly using Snippy v4.6.0 [49] and confirming that 0 variants were called. We used (i) Prokka v1.14.6 [50] and (ii) RATT v1.0.3 [51] to (i) annotate our reference strains using a *P. aeruginosa* pan-genome database collated by Bactopia, and to (ii) transfer gene annotations from PAO1 to their respective positions in each of the reference strains, respectively.

### Variant calling

We used Snippy v4.6.0 (39) to call variants from the other 296 isolates against their respective reference strain and create a core genome alignment. Using PhyML v3.3.20211231 (43), we created a maximum likelihood phylogeny. Then, using VCFtools v0.1.16 (44) and Disty McMatrixface v0.1.0 (45), we generated a pairwise SNP matrix for each patient. For Disty, we only considered alleles in the core genome and chose to ignore ambiguous bases pairwisely (-s 0). We then employed SnpEff and SnpSift v4.3t (46) to identify the affected genes and sort the variants by predicted effect. We identified hypermutators in these populations by the presence of non-synonymous mutations in *mutL*, *mutS*, and *uvrD* [52].

### Antimicrobial susceptibility testing

To assess antimicrobial susceptibility profiles, we followed the guidelines and standards provided by the Clinical and Laboratory Standards Institute (CLSI) *Performance Standards for Antimicrobial Susceptibility Testing M100S*, 30^th^ edition. We first grew all isolates overnight in LB in 24-well microtiter plates at 37℃ with shaking at 200 rpm. We diluted cultures to a Macfarland standard of 0.5 (OD_600_ ∼0.06) and streaked a lawn on 100×15 mm Petri dishes with 20 ml Mueller-Hinton agar using pre-sterilized cotton swabs. We then stamped amikacin (AK), meropenem (MEM), piperacilin-tazobactam (TZP), ciprofloxacin (CIP), tobramycin (TOB), and ceftazidime (CAZ) on each plate and incubated for 17 h at 37℃. We measured the zone of inhibition (ZOI) at 17 h and classified the values as resistant, intermediate, or susceptible per the established CLSI interpretive criteria. We used *P. aeruginosa* strain ATCC 27853 as a quality control. We tested all isolates in biological triplicates. We ran a Mann-Whitney U test to compare the means of antimicrobial susceptibilities between hypermutators and normomutators (non-hypermutators) and a Pearson’s correlation coefficient to determine relationships between susceptibilities to different antimicrobials, both using ⍺ = .05.

### Principal components analysis

We conducted a principal components analysis of the antimicrobial susceptibility data in R v4.3.0 using a singular value decomposition approach.

### Resistome genotyping

We assessed genotypes relevant to resistance by uploading the *de novo* assemblies to the Resistance Gene Identifier (RGI) v6.1.0 web portal, which predicts resistomes using the Comprehensive Antibiotic Resistance Database (CARD) v3.2.6 [53]. We excluded loose and nudge hits from this analysis.

### Enrichment analysis

We conducted an enrichment analysis to determine which functional categories of genes were differentially impacted by mutations than would be expected by random chance. We used an in-house Python script to retrieve the PseudoCAP functional group of each gene where a non-synonymous SNP or microindel was identified. We accounted for the varying lengths of genes in each functional category in our analysis, based off their lengths and prevalence in the PAO1 genome. We used a chi-squared goodness of fit test to conduct the enrichment analyses for Patients 1-3 to determine which functional categories were disproportionately impacted by non-synonymous variants. We used the R package XNomial v1.0.4 [54] to conduct an exact multinomial goodness of fit test using Monte-Carlo simulations for Patient 4 because the SNP frequencies of Patient 4 did not meet the assumptions for a chi-squared test. Given the formula for calculating the chi-squared statistic: 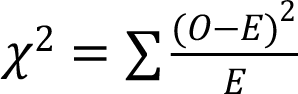, if the 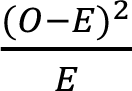 value for a particular PseudoCAP functional category was in the top 30 percentile of all) values (top 8 of 27 total categories) in the analyses of at least three patients, we noted this as an enrichment.

### Predicting putative recombination events

We input the core genome alignment from each patient to Gubbins v3.3.0 [55] to predict potential recombinant regions in each population.

### Analyzing growth curves

To assess growth, we cultured strains for 24 h in 96-well microtiter plates (Corning) at 37℃ static, in 200μL synthetic cystic fibrosis sputum medium (SCFM) [42], shaking for 4 s before reading optical density at 600 nm every 20 min. We tested all clinical isolates in biological triplicates. We used GrowthCurver [56] to analyze the resulting growth curves and calculate growth rate (r). We then assessed the relationship between growth rate and susceptibility profiles using a linear mixed model in brms [57].

### Visualizations

We conducted graphical analyses in R v4.3.0.

### Data availability

The sequences in this study will be made available in the NCBI SRA database upon publication.

## Results

### Description of the four patient cohort selected for this study

The four individuals selected for this study were aged 24-31 years and had been chronically infected with *P. aeruginosa* for 10-15 years at the time of sampling. All four individuals had at least one copy of the F508del CFTR mutation, but none were on CFTR modulator therapy. Patients 1, 2, and 4 were seeking outpatient treatment for an acute pulmonary exacerbation at the time of sampling, while Patient 3 was in stable medical condition. These individuals were in the early (%FEV_1_ > 70) to intermediate (%FEV_1_ ≤ 70, ≥ 40) stages of lung disease, with %FEV_1_ scores ranging from 60.30% to 74.92%. The antibiotic regimens for each patient at the time of sampling were as follows: Patient 1 was receiving inhaled tobramycin and oral azithromycin; Patient 2 was receiving inhaled tobramycin and oral trimethoprim/ sulfamethoxazole; Patient 3 was receiving inhaled tobramycin, oral azithromycin, and inhaled aztreonam; and Patient 4 was receiving inhaled tobramycin, oral trimethoprim/ sulfamethoxazole, and oral levofloxacin (**Table 1**).

**Table 1.** Metadata on the four patients in our cohort: sex, cystic fibrosis transmembrane conductance regulator (CFTR) mutation status, length of *P. aeruginosa* infection, clinical status, forced expiratory volume (% FEV1), modulator therapy, antibiotic treatment at time of sampling, and dominant infection strain type.

|  | Patient 1 | Patient 2 | Patient 3 | Patient 4 |
| --- | --- | --- | --- | --- |
| **Patient Sex** | F | F | F | M |
| **CFTR Mutation** | F508del/R1162X | F508del/F508del | F508del/L467P | F508del/ 621+1G->T |
| **Length of *Pa* infection** | 15 years, 2 months | 12 years, 5 months | 10 years, 4 months | 13 years |
| **Clinical status** | APE Outpatient | APE Outpatient | Stable | APE Outpatient |
| **FEV<sub>1</sub> (%)** | 67.96% | 74.92% | 67.83% | 60.30% |
| **Modulator Therapy** | None | None | None | None |
| **Antibiotic Treatment** | Inhaled tobramycin, oral azithromycin | Inhaled tobramycin, oral Trimethoprim / Sulfamethoxazole | Inhaled tobramycin, inhaled aztreonam, oral azithromycin | Inhaled tobramycin, oral Trimethoprim / Sulfamethoxazole, oral levofloxacin |
| **Dominant ST** | 870 | 2999 | 1197 | 274 |

### *P. aeruginosa* populations display significant within-patient diversity in antimicrobial resistance profiles

In order to assess diversity in AMR, we selected 75 isolates from a single sputum sample of each of the four individuals for a total of 300 isolates. Using a standard disc diffusion assay, we assessed these 300 isolates for their susceptibilities to six antimicrobials commonly prescribed in CF treatment: amikacin, meropenem, piperacilin-tazobactam, ciprofloxacin, tobramycin, and ceftazidime (**Tables S1-S4**). Zone of inhibition values within a population for a given antibiotic displayed a statistical range (minimum subtracted from the maximum value of a population) between 6 and 25.3 mm, with an average of 12.75 mm. Standard deviations of these values ranged from 1.4 to 8.0 mm, with an average standard deviation of 3.0 mm. The majority of isolates presented values well within the range of susceptibility for the tested antibiotics, despite ineffective clearing of infection in the clinic for these patients chronically infected with *P. aeruginosa* (**Fig. 1**). Only two patients harbored isolates that tested in the range of clinical resistance to any antimicrobial: amikacin, ciprofloxacin, and tobramycin for Patient 1; and ciprofloxacin for Patient 3. Three of the four patients harbored isolates that presented phenotypes spanning across the clinical thresholds for resistant, intermediate, and susceptible for at least one, if not multiple, antibiotics. Principal components analysis of these values show that isolate antimicrobial sensitivity phenotypes cluster by patient (**Fig. 2**).

**Figure 1.**
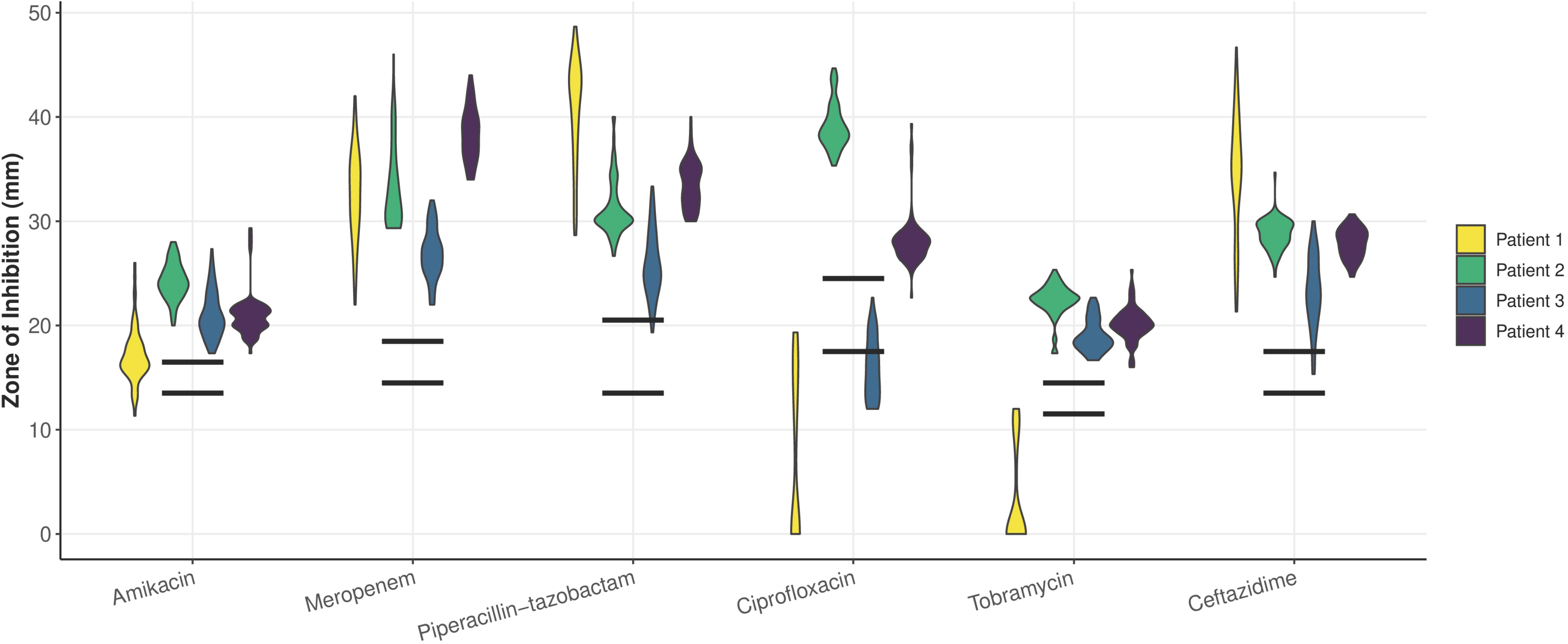
Violin plot of the antimicrobial susceptibility profiles of all four populations against amikacin, meropenem, piperacillin-tazobactam, ciprofloxacin, tobramycin, and ceftazidime as measured by zone of inhibition in a standard disc diffusion assay shows phenotypic diversity across all populations. Data points are clustered and colored by respective patient, with each individual violin plot representing 75 isolates from a single patient. Black horizontal bars indicate the cut-off values for susceptibility (top bar) and resistance (bottom bar) for each antibiotic as determined by the Clinical and Laboratory Standards Institute (CLSI). Clinical thresholds for resistance to amikacin, meropenem, piperacillin-tazobactam, ciprofloxacin, tobramycin, and ceftazidime are 14, 15, 14, 18, 12, and 14 mm, respectively. Clinical thresholds for sensitivity to these antimicrobials are 17, 19, 21, 25, 15, and 18 mm, respectively.

**Figure 2.**
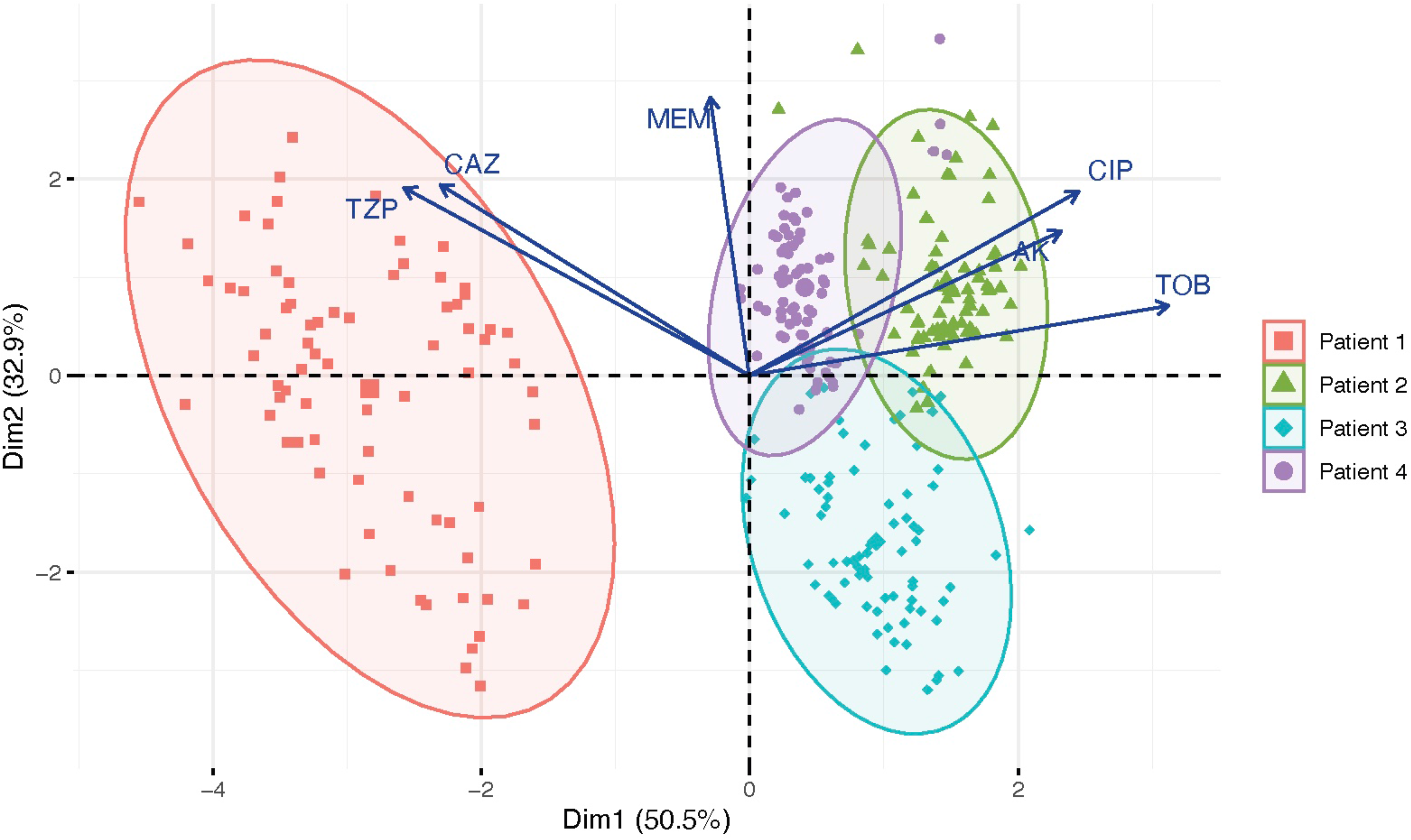
Principal components analysis plot of antimicrobial sensitivities shows that isolates cluster by patient. 50.5% of the variance in antimicrobial sensitivities is demonstrated by dimension 1, and 32.9% of the variance is demonstrated by dimension 2. Vectors demonstrate to what degree each variable (i.e., antimicrobial) influences the principal components.

### The four patients are chronically infected by a single *P. aeruginosa* strain, populations of which display a range of genomic diversity levels

In order to quantify the level of within-patient genomic diversity for these populations, we sequenced the 75 isolates from each of the four individuals of this cohort. We prepared the sequences of all 300 isolates using *de novo* assembly and annotation. We assembled the genomes in 20 to 444 contigs (mean = 53 contigs; **Table S5**). Genomes in this dataset ranged in size from 5,888,197 to 6,746,489 nucleotides, with 5,209 to 5,970 genes (**Table S5**). The median genome sizes of isolates sourced from Patients 1-4, respectively, were 6,222,786, 6,331,110, 6,742,689, and 6,308,671 nucleotides, with 5,523, 5,571, 5,964, and 5,567 genes, respectively (**Table S5**). A phylogenetic tree of the core genome alignment revealed that the populations infecting Patients 1, 2, and 4 clustered closely with PAO1, while that of Patient 3 more closely resembled PA14 (**Fig. S1**). Strain typing of the isolates showed that there was a single *P. aeruginosa* strain type in each patient— ST870, ST2999, ST1197, and ST274 for Patients 1-4, respectively (**Table 1**). For the rest of the text, we will simply refer to each population by its respective patient number.

We assessed the genomic diversity in these populations according to the number of single nucleotide polymorphisms (SNPs) and microindels (insertions and deletions). We found that genomic diversity varied significantly between patients. The total number of unique SNPs discovered across 75 isolates for Patient 1 was 4,592 (maximum number of pairwise SNPs = 611, median number of pairwise SNPs = 199, mean = 208); for Patient 2 was 1,972 (max. = 326, median = 145, mean = 118); for Patient 3 was 1,638 (max. = 150, median = 76, mean = 87); and for Patient 4 was 31 (max. = 8, median = 1, mean = 3) (**Fig. 3**; **Table 2**). Across the population of Patient 1 we found 498 unique microindels, 307 for Patient 2, 330 for Patient 3, and 14 for Patient 4 (**Table 2**).

**Figure 3.**
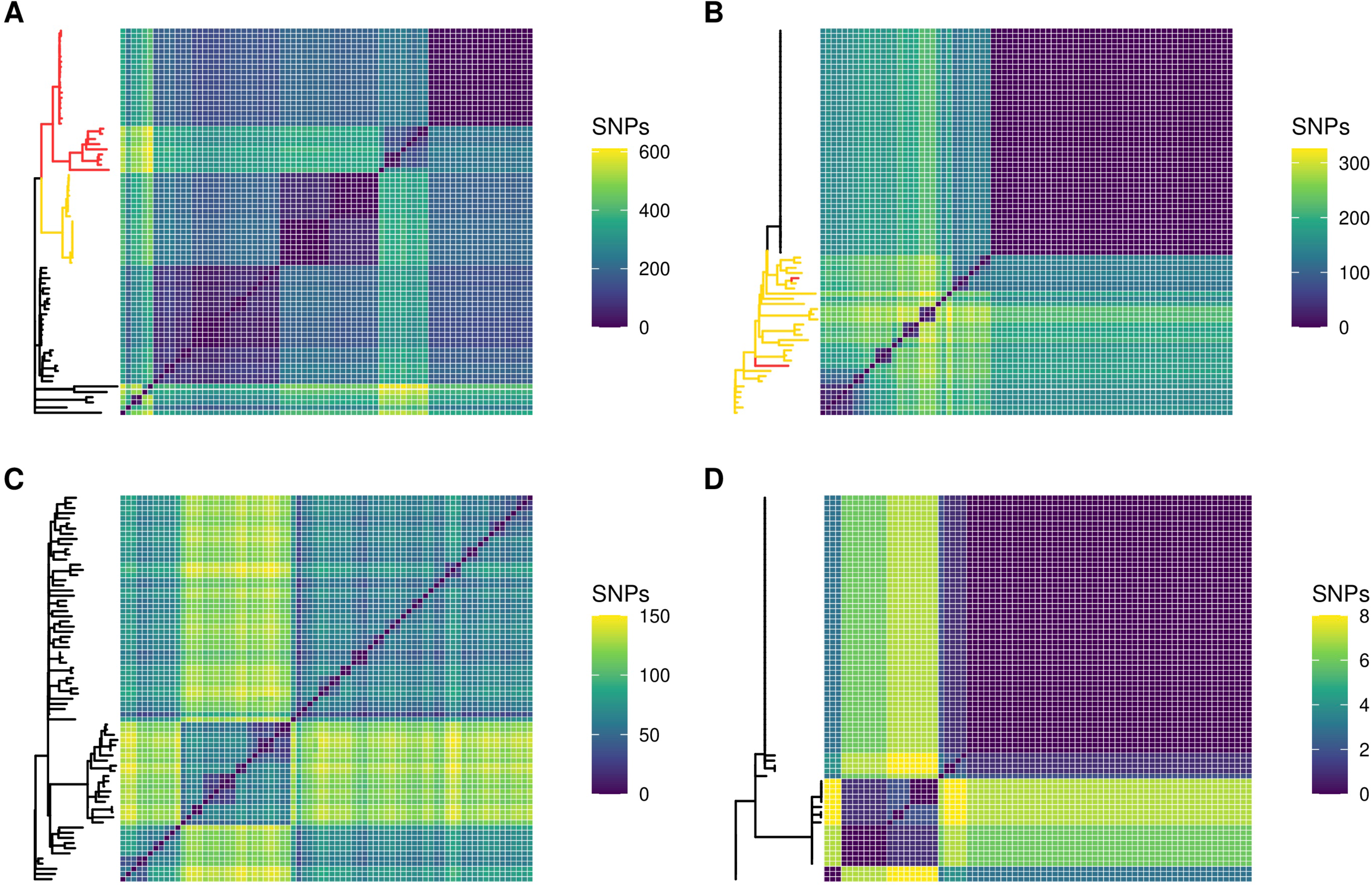
Genomic diversity as measured by core genome SNPs varies greatly from one population to another. Populations are presented in order of decreasing genomic diversity: Patient 1 (A), Patient 2 (B), Patient 3 (C), and Patient 4 (D). Each matrix represents the pairwise comparison of SNPs across all 75 isolates within a population against each other, and each population is composed of a single strain type. Isolates with one DNA mismatch repair mutation are highlighted in yellow on phylogenies. Isolates with two DNA mismatch repair mutations are highlighted in red.

**Table 2.** Genetic variations in each population: single nucleotide polymorphisms (SNPs), multiple nucleotide polymorphisms (MNPs), and insertions and deletions (indels).

|  | Patient 1 | Patient 2 | Patient 3 | Patient 4 |
| --- | --- | --- | --- | --- |
| **Total # unique SNPs/ MNPs** | 4592 | 1972 | 1638 | 31 |
| **# SNPs/ MNPs separating most divergent isolates** | 611 | 326 | 150 | 8 |
| **Non-synonymous SNPs/ MNPs** | 2803 | 1294 | 1024 | 24 |
| **Synonymous SNPs/ MNPs** | 1248 | 484 | 425 | 5 |
| **SNPs in non-coding regions** | 541 | 194 | 189 | 2 |
| **Total # indels** | 498 | 307 | 330 | 14 |
| **Indels in non-coding regions** | 204 | 99 | 115 | 2 |

### Genomic diversity may not be a consistent predictor of antimicrobial resistance diversity in a population

We next determined whether genomic diversity could serve as a predictor of diversity in AMR phenotypes in our cohort. We hypothesized that genetically diverse populations would also display more diversity in AMR. We chose to quantify genomic diversity in terms of SNPs. We quantified AMR diversity using the number of distinct AMR profiles (i.e., distinct zone of inhibition values) for a given antibiotic within a population. Total SNP count in a population was a strong indicator of AMR diversity for amikacin (R^2^ = .90, F(1, 2) = 18.94, *p* = .049), meropenem (R^2^ = .93, F(1, 2) = 25.3, *p* = .037), and piperacilin-tazobactam (R^2^ = .95, F(1, 2) = 39.86, *p* = .024). However, SNP count was a poor indicator of AMR diversity for ciprofloxacin (R^2^ = .12, F(1,2) = .27, *p* = .65) and ceftazidime (R^2^ = .71, F(1,2) = 4.78, *p* = .16), and was inversely related to AMR diversity for tobramycin (R^2^ = .97, F(1,2) = 66.61, *p* = .015) (**Fig. S2**). We next used the number of distinct CARD resistance genotype profiles within a population (**Fig. 4**) as a proxy for genomic diversity to eliminate bias from SNPs not relevant to AMR and to account for the epistatic or synergistic effect that combinations of various alleles may have. This yielded similar results to the previous analysis (**Table S6**). We then instead used the standard deviation of zone of inhibition values within a population as a proxy for AMR diversity to see if this would improve the strength of the association between genomic diversity and phenotypic diversity for these antimicrobials. We found that the number of distinct CARD profiles within a population was a better predictor of standard deviation for ciprofloxacin (R^2^ = .79, F(1,2) = 7.35, *p* = .11), tobramycin (R^2^ = .77, F(1,2) = 6.73, *p* = .12), and ceftazidime (R^2^ = .81, F(1,2) = 8.44, *p* = .10), though these associations were still not significant (**Fig. S3**).

**Figure 4.**
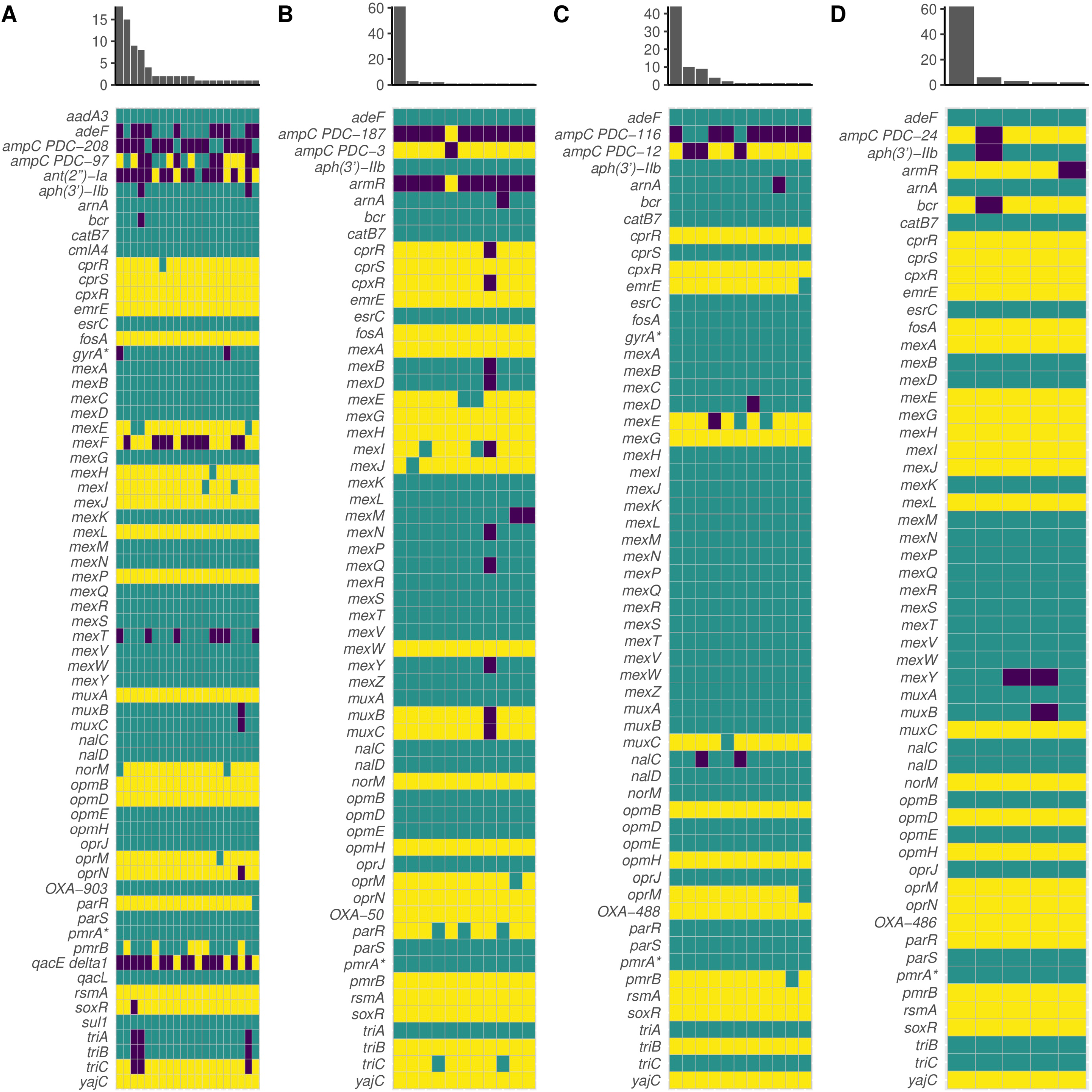
Visualized resistomes of Patients 1 (A), 2 (B), 3 (C), and 4 (D) as predicted by the Comprehensive Antibiotic Resistance Database Resistance Gene Identifier (CARD RGI) demonstrate decreasing levels of resistome diversity. Yellow indicates a perfect hit to the database, teal indicates a strict hit, and purple indicates no hit (or loose hit in some cases). X-axis of the histogram indicates the number of unique resistome profiles in the population, and y-axis indicates the number of isolates in the population that share a unique resistome profile. An asterisk (*) indicates a gene with resistance conferred by a mutation (i.e. CARD RGI protein variant model).

### *P. aeruginosa* diversity is primarily driven by *de novo* mutations, especially mutations in DNA mismatch repair

We next wanted to further understand the processes by which *P. aeruginosa* diversified in our cohort. We first sought to predict putative recombination events. In Patients 1-4, 527 (11.5%), 19 (<1%), 86 (5.25%), and 0 SNPs were predicted to be in 31, 3, 17, and 0 recombinant regions, respectively. These data show that *de novo* mutation was a much more prominent driver of intra-specific diversity than recombination in our particular cohort. As expected, we found that the infections with the highest SNP diversity harbored strains with DNA MMR mutations. Patients 1 and 2 harbored DNA MMR mutants (hypermutators); however, we found no hypermutators in Patients 3 or 4 (**Fig. 3**). The phylogeny of Patient 1 indicates that a non-synonymous SNP in *mutS* (Ser31Gly) evolved first in the population, after which a frameshift deletion in *mutS* (Ser544fs) piggybacked. In total, *mutS* mutants comprise 61.3% of this population. In Patient 2, a non-synonymous SNP in *mutL* resulting in a pre-mature stop codon (Glu101*) evolved first, found in 41.3% of the population. Two of these *mutL* mutants further independently acquired a single non-synonymous mutation in *mutS* (Phe445Leu, Ala507Thr) (**Fig. 3**).

In Patient 1, there were two distinct branches of the phylogenetic tree, one with hypermutators and the other composed of normomutators (38.7%) (**Fig. 3**). Interestingly, there was a significant amount of genetic diversity within both the normomutators (mean SNP distance = 156.9 SNPs, median = 91 SNPs) and hypermutators (mean = 174.6 SNPs, median = 197 SNPs). There was a distinct small cluster of normomutator isolates that significantly diverged from the others. Of the hypermutators, these further diverged into those with one DNA MMR mutation (39.1%) and those with two MMR mutations (60.9%). In Patient 2, there was largely a lack of genetic diversity in the normomutators (mean = 0.36 SNPs, median = 0 SNPs), with one clone dominating 48% of the population (**Fig. 3**). The emergence of hypermutators appears to have been responsible for the large majority of all the genetic diversity in this population (mean = 211.2 SNPs, median = 224 SNPs). In Patient 3, there were three major lineages, comprising 58.7%, 26.7%, and 14.7% of the total population (mean = 61.9, 55.5, and 65.4 SNPs; median = 62, 61, and 64 SNPs, respectively; **Fig. 3**). In Patient 4, there was one dominant clone encompassing 66.6% of the population, with a small number of SNPs (mean = 4 SNPs, median = 3 SNPs) differentiating the other 33.3% of the population (**Fig. 3**).

### Hypermutation can drive the evolution of increased susceptibility to antimicrobials, even under apparent selective pressure

As our cohort had two populations with DNA MMR mutants, we used this opportunity to ascertain how hypermutation drives the evolution of AMR. In Patient 1, AMR genotypes cluster by DNA MMR genotype. Hypermutators were significantly more resistant to amikacin than normomutators (U = 315.5, *p* = .00013) (**Fig. 5**), although this difference could not be attributed to any hits in the CARD database. Hypermutators were also significantly more resistant to beta-lactams piperacilin-tazobactam (U = 457.5, *p* = .023) and ceftazidime (U = 428, *p* = .0095), although there was no significant difference in the resistance profiles of hyper-and normomutators with regards to the beta-lactam meropenem (U = 630, *p* =.69) (**Fig. 5**). Some normomutators in this population acquired a SNP in *ampC* (461 A > G, Asp154Gly) (**Fig. 4**), which was associated with increased sensitivity to piperacilin-tazobactam (U = 320, *p* = .0014) and ceftazidime (U = 342.5, *p* = .0034). Of the isolates with one DNA MMR mutation, some lost *ampC* entirely, also associated with increased susceptibility to ceftazidime (U = 106, *p* = .0019). Of the isolates with both DNA MMR mutations, some had acquired a SNP in *ampC* (1066 G > A, Val356Ile), which appeared to increase their resistance to piperacilin-tazobactam (U = 12, *p* < .00001) and ceftazidime (U = 8, *p* < .00001) (**Fig. 4**).

**Figure 5.**
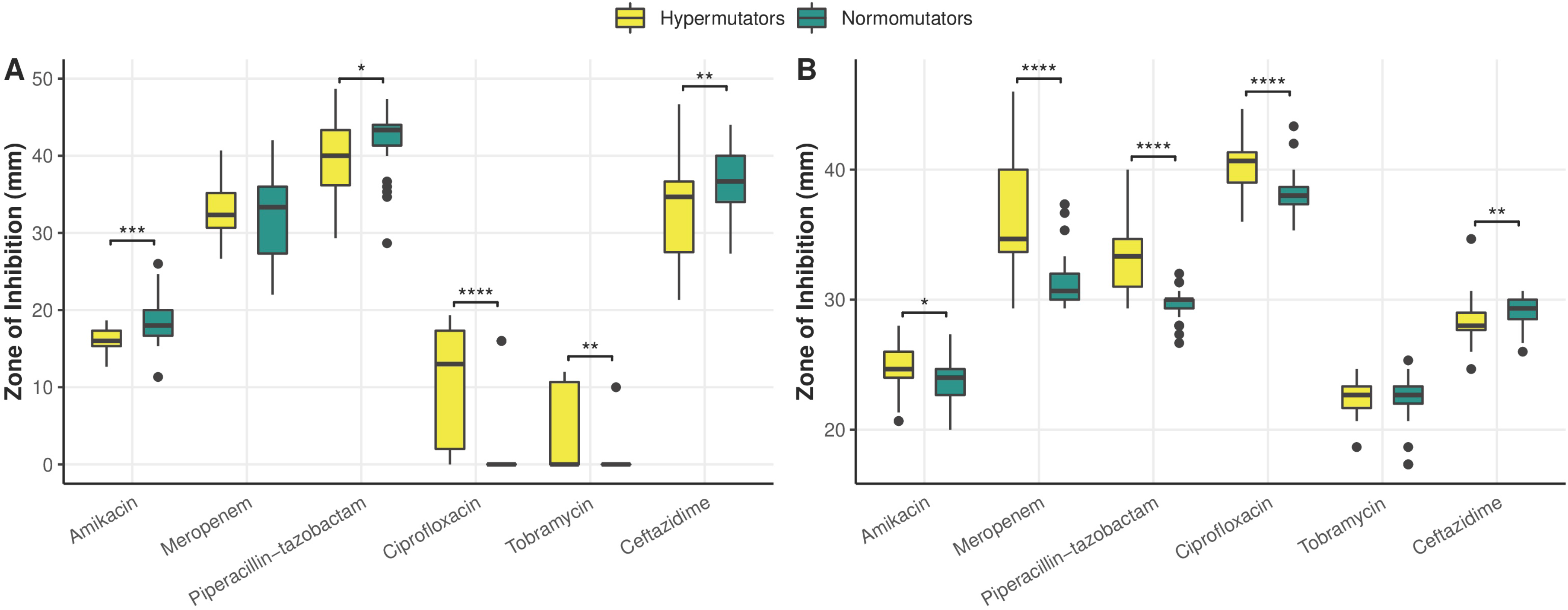
Comparative antimicrobial susceptibility profiles of hypermutators and normomutators in Patient 1 (A) and Patient 2 (B) as measured by zone of inhibition in a standard disc diffusion assay highlight increased sensitivities and resistance levels by hypermutators. (A) In Patient 1, hypermutators were significantly more resistant to amikacin (U = 315.5, p = .00013), piperacilin-tazobactam (U = 457.5, p = .023), and ceftazidime (U = 428, p = .0095) than normomutators, although there was no significant difference in the resistance profiles of hyper-and normomutators in regards to meropenem (U = 630, p =.69). Hypermutator isolates in Patient 1 displayed zone of inhibition (ZOI) values that were on average 10 times larger for ciprofloxacin (U = 218, p < .00001) and >13 times larger for tobramycin (U = 379.5, p = .0018) than normomutators, and isolates with both DNA MMR mutations in this population additionally presented ZOI values that were 36 times larger than normomutators for tobramycin (U = 172.5, p < .00001), indicating increased sensitivity displayed by hypermutators. (B) In Patient 2, hypermutators displayed increased susceptibility to amikacin (U = 479, p = .029), meropenem (U = 194, p < .00001), piperacilin-tazobactam (U = 121.5, p < .00001), and ciprofloxacin (U = 213.5, p < .00001) relative to normomutators. Hypermutators in Patient 2 were more resistant to ceftazidime (U = 417.5, p = .0045). There was no statistically significant difference between the tobramycin susceptibility profiles of hyper- and normomutators in this population (U = 634.5, p = .61). (*) indicates p ≤ .05, (**) indicates p ≤ .01, (***) indicates p ≤ .001, and (****) indicates p < .0001 in a Mann-Whitney U test. Clinical thresholds for resistance to amikacin, meropenem, piperacillin-tazobactam, ciprofloxacin, tobramycin, and ceftazidime as determined by the CLSI are 14, 15, 14, 18, 12, and 14 mm, respectively. Clinical thresholds for sensitivity to these antimicrobials are 17, 19, 21, 25, 15, and 18 mm, respectively.

Interestingly, hypermutator isolates in this population displayed zone of inhibition values that were on average 10 times larger for ciprofloxacin (U = 218, *p* < .00001) and >13 times larger for tobramycin (U = 379.5, *p* = .0018) than normomutators, indicating increased sensitivity of hypermutators to these antimicrobials (**Fig. 5**). Isolates with both DNA MMR mutations in this population additionally presented ZOI values that were 36 times larger than normomutators for tobramycin (U = 172.5, *p* < .00001) (**Fig. 5**). The altered ciprofloxacin phenotype may be explained in part by SNPs in *gyrA* (248 T > C, Ile83Thr) or *norM* (61 G > A, Ala21Thr) (U = 38.5, *p* < .00001) (**Fig. 4**). However, there were isolates in this population whose phenotypes were not ostensibly explained by either of these genotypes. The increased susceptibility to tobramycin was strongly linked to the aforementioned SNP in *norM* (U = 31.5, *p* < .00001) (**Fig. 4**). We observed apparent evidence of one of these hypermutators reversing this increased susceptibility to tobramycin by acquisition of the aminoglycoside nucleotidyltransferase *ant(2”)-Ia* (**Fig. 4**). There was additionally a normomutator isolate with an outlier tobramycin susceptibility phenotype. Interestingly, 12 isolates from Patient 1 had improved growth in the presence of tobramycin (determined by visual observation of denser growth in the region surrounding the antibiotic disc in a disc diffusion assay), a phenotype which could not be explained by any hits in the database. All of the normomutator isolates had a truncated *mexF* (**Fig. 4**), although this did not appear to impact any of the tested phenotypes.

In Patient 2, hypermutators displayed increased sensitivities to meropenem (U = 194, p < .00001), piperacilin-tazobactam (U = 121.5, *p* < .00001), and ciprofloxacin (U = 213.5, *p* < .00001) relative to normomutators (**Fig. 5**). This appeared to be caused in part by a SNP in *mexB* (2257 T > C, Trp753Arg) shared by all hypermutators in this population. However, there were outliers whose phenotype could not be explained by this genotype. Hypermutators were also more susceptible to amikacin (U = 479, *p* = .029) and more resistant to ceftazidime (U = 417.5, *p* = .0045) (**Fig. 5**), although these strains harbored no apparent genes or SNPs associated with these phenotypes in the CARD database. There was no statistically significant difference between the tobramycin susceptibility profiles of hyper- and normomutators in this population (U = 634.5, *p* = .61) (**Fig. 5**). One hypermutator isolate in Patient 2 had an unusual density of truncated pseudogenes, 10 of which are involved in resistance mechanisms and 9 of which specifically play roles in resistance-nodulation-cell division efflux— *mexY*, *mexQ*, *mexN*, *cpxR*, *muxB*, *muxC*, *mexI*, *mexB*, *mexD*, and *cprR* (**Fig. 4**). Although RGI denoted these genes as missing due to truncation, this isolate was equally or more resistant to every antimicrobial tested relative to other DNA MMR mutants in the population, suggesting that many of these genes were still functional.

In the two normomutator populations, there was significantly decreased resistome diversity. In Patient 3, a SNP in *ampC* (716 T > C, Val239Ala) was associated with increased resistance to ceftazidime (U = 165.5, *p* < .00001) and piperacillin-tazobactam (U = 312.5, *p* = .0045) (**Fig. 4**). Some of the isolates with this SNP additionally were missing *nalC* (**Fig. 4**) and displayed increased susceptibility to meropenem (U = 172.5, *p* = .01778) relative to other isolates. In Patient 4, a truncation in *mexY* was strongly linked to variations in sensitivities to amikacin (U = 35, *p* = .0031), piperacillin-tazobactam (U = 22.5, *p* = .0012), ciprofloxacin (U = 0, *p* = .0002), and tobramycin (U = .5, *p* = .00022) (**Fig. 4**). Surprisingly, isolates missing a hit to *aph(3’)-IIb* were more resistant to aminoglycosides amikacin (U = 11.5, *p* = .00014) and tobramycin (U = 55, *p* = .00308), and those missing a hit to *ampC* were more resistant to ceftazidime (U = 62, *p* = .0048) (**Fig. 4**). Seeing as these relationships are unexpected, it is likely that there are other genetic variations not cataloged in the CARD database, or epistatic interactions, that are influencing these phenotypes.

### Protein export/ secretion systems and transcriptional regulators are hotspots for *de novo* mutations in these populations

To determine whether these populations were enriched for mutations in genes with roles in resistance, we categorized non-synonymous SNPs and microindels that occurred within coding regions of genes according to the PseudoCAP functional categories and conducted an enrichment analysis. We did not find that AMR genes were enriched for such variants in this cohort (**Fig. S4**). However, we found that protein secretion and export apparatuses and transcriptional regulators were enriched for such mutations (**Fig. S4**). Additionally, two of the four genes impacted by non-synonymous mutations in all four populations in this study were related to protein secretion, *fha1* and *pscP* (**Table S7**). We found that phage/transposon/plasmid genes were less likely to be impacted by such mutations (**Fig. S4**).

Non-coding RNAs were also less likely to be impacted by mutations than other functional categories (**Fig. S4**; see **Table S8** for all supporting statistical values), which is unsurprising given that small non-coding RNAs are known to hold important regulatory functions in bacteria [58]. 57 genes were impacted by non-synonymous mutations in at least 3 of 4 patients, which included genes with previously described functions in alginate biosynthesis, primary metabolism, antibiotic resistance and efflux, iron uptake, biofilm formation, stress response, amino acid biosynthesis, type IV pili, lipopolysaccharide, quorum sensing, and virulence (**Table S9**). A full list of all SNPs discovered in this dataset can be found in **Tables S10-S13**.

### Populations display poor evidence for evolutionary trade-offs to explain heterogeneity in resistance profiles

We next wanted to ascertain if there was any evidence of evolutionary trade-offs involving AMR in these populations. Collateral sensitivity is sensitive to genetic background [17, 19, 59, 60] and must be proven robust across a wide range of genetic backgrounds in order to be broadly applicable as a therapeutic strategy [61]. Therefore, we searched for evidence of collateral sensitivity within our populations, and additionally for evidence of trade-offs between AMR and fitness (i.e., growth rate) in a CF sputum-like medium, SCFM [42]. Using the Pearson’s correlation coefficient, we found no evidence of collateral sensitivity across any of the six antimicrobials tested for any patient (**Fig. 6**). A principal components analysis conducted for each patient further confirmed this, and showed that cross-resistance and cross-sensitivity patterns differed between patients (**Fig. S5**). We analyzed growth curves for all 300 isolates (**Tables S14-S17**) and using a linear mixed model, determined that there was not a significant relationship between resistance and fitness for any of the tested antimicrobials (**Fig. S6; Table S18** for supporting code and statistical values).

**Figure 6.**
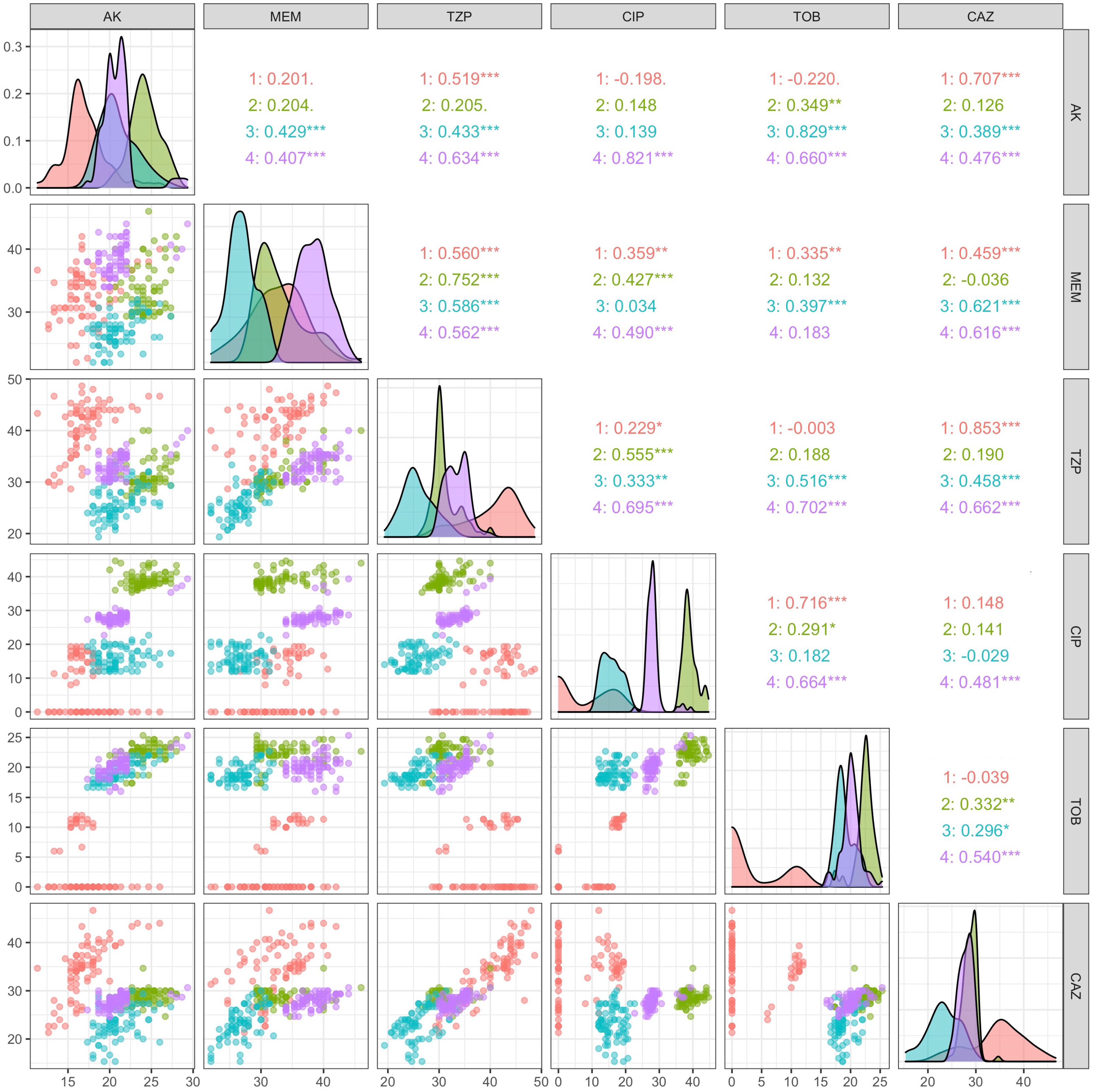
Lack of statistically significant negative correlations between any two antimicrobial susceptibility profiles in a Pearson’s correlation provides no evidence for collateral sensitivity trade-offs. Pearson’s correlation coefficient (upper right quadrant), scatterplots (lower left quadrant), and density plots (diagonal) for pairwise comparisons of susceptibility profiles across all six tested antimicrobials: amikacin (AK), meropenem (MEM), piperacillin-tazobactam (TZP), ciprofloxacin (CIP), tobramycin (TOB), and ceftazidime (CAZ).

## Discussion

The goal of this project was to better understand how genomic diversification in *P. aeruginosa* CF lung populations drives the evolution of AMR. For this study, we selected four distinct patients with varying levels of *P. aeruginosa* genomic population diversity, ranging from a few dozen to multiple thousands of SNPs within a given population. We found that (i) genomic diversity was not consistently a reliable predictor of AMR diversity for this cohort; (ii) hypermutators in one population evolved increased sensitivity to tobramycin, even when undergoing treatment by tobramycin; and that (iii) there was no evidence for collateral sensitivity or trade-offs between AMR and fitness in these populations.

Previous studies have reported both on genomic and phenotypic diversity of *P. aeruginosa* in CF airways [6, 34–40]; however, the clinical implications of genomic diversity within these populations on resistance diversity have not been fully assessed. Our results suggest that genomic diversity may not be a reliable predictor of phenotypic diversity for all antibiotics. However, there are a number of limitations to this finding: (i) our sample size for this analysis was small; (ii) we cannot account for diverse genotypes that result in converging phenotypes; and (iii) there are likely many genetic variants that act on AMR that have not been catalogued in CARD. Nonetheless, we highlight that Patient 4 displayed a number of distinct AMR profiles that was, in the case of ciprofloxacin, comparable to that of Patient 1, which had 148x more SNPs and 4x as many distinct CARD genotype profiles within the population. In the case of tobramycin, Patient 4 displayed more distinct AMR profiles and higher zone of inhibition standard deviation values compared to Patients 2 and 3, which both had 2x as many distinct CARD genotype profiles and over 53x more population SNPs compared to Patient 4. Ultimately, because of our limited ability at present to predict the phenotypic impact of novel genetic variants or the epistatic interactions of alleles *in silico*, it may prove challenging to ascertain the phenotypic heterogeneity of an infection in a parsimonious manner that could be translated to the clinic [41]. In addition to improved *in silico* capabilities, greater understanding of the social interactions that impact how co-infecting microbes with varying resistance levels collectively respond to antibiotic treatment and development of reliable methodology for assessing population-level resistance are also necessary. Considering the impact of polymicrobial interactions has certainly been shown to add an additional layer of complexity in predicting the antimicrobial sensitivity profiles of diverse infections [32, 62], although there is still uncertainty in the degree to which various species of pathogens spatially co-exist and interact in the CF lung. Improved understanding of how these social dynamics influence AMR may be instrumental in future approaches for tackling chronic infections.

Our data further highlight that even our ability to assess resistance at the isolate-level is inadequate. Though the majority of the isolates selected for this study demonstrated sensitivity to nearly every antibiotic *in vitro*, these testing results likely underestimate resistance levels *in situ*, given that these populations have persisted within the lung for over a decade and that only one population displayed clinical resistance to tobramycin, despite all four individuals in this cohort undergoing treatment with inhaled tobramycin. These findings are in accordance with the wide array of literature that has already called into question the utility of antimicrobial susceptibility testing in the clinic, which falls short in reproducing the hypoxic CF microenvironment and the biofilm mode of growth displayed by *P. aeruginosa* in this biological context, and ultimately fails in predicting patient outcomes [5, 63, 64]. Still, we found it particularly unusual that two of our populations did not display clinical resistance to any of the antimicrobials tested *in vitro*, as prior studies on AMR diversity of *P. aeruginosa* in CF lungs have generally demonstrated high prevalence of *in vitro* resistance within populations [34–38].

Two limitations of our study are that we were unable to obtain full treatment histories for these patients, and that the pre-selected panel of antimicrobials tested did not include all those that the four patients were undergoing treatment with at the time of sampling (i.e., aztreonam, azithromycin, trimethoprim-sulfamethoxazole, and levofloxacin). Disc diffusion data on these antimicrobials in addition to treatment histories of these patients could potentially illuminate the reasons for treatment failure and explain the presence of strains resistant to amikacin and ciprofloxacin. However, (i) the mechanisms of resistance for levofloxacin and aztreonam closely overlap with those of the other aminoglycoside and beta-lactam antibiotics tested; (ii) trimethoprim-sulfamethoxazole is not prescribed as a treatment for *P. aeruginosa*; and (iii) azithromycin does not display conventional antimicrobial activity against *P. aeruginosa*, but rather, inhibits quorum sensing (therefore, rendering traditional disc diffusion testing of this drug non-viable). Therefore, we believe that our results still broadly provide coverage of the spectrum of relevant antimicrobial sensitivities displayed by these populations. We were additionally concerned to discover strains with increased growth in the presence of tobramycin, as inhaled tobramycin is one of the most commonly prescribed drugs for CF patients with *P. aeruginosa* infection. It may be that tobramycin is being catabolized by these strains to aid in growth, although further investigation is needed to test this hypothesis.

Combining single-isolate whole genome sequencing and phenotypic characterization approaches further allowed us to understand how the evolution of genotypes and combination of alleles impact AMR within a population. Although we were able to identify a number of candidate genotypes responsible for these phenotypic variations, there were a number of unexplained phenotypic outliers, highlighting the presence of novel genetic signatures of AMR or allelic interactions influencing AMR phenotype. Previous reports have primarily focused on the role that hypermutation plays in evolving increased AMR in clinical *P. aeruginosa* populations [65–71]. We found ample evidence that hypermutation can also lead to increased susceptibility, such as the hypermutator isolates in Patient 1 that were significantly more sensitive to tobramycin, despite this patient undergoing treatment with inhaled tobramycin. This may be a function of antimicrobial treatment regimens exerting uneven selection pressure on the population. Or, it may be that the evolution of genetic resistance for these populations is inconsequential because antimicrobials are failing to penetrate phenotypic barriers, such as biofilms, and other mechanisms of antibiotic tolerance, including persister cells with reduced metabolic activity in the microaerophilic lung [72–78]. Although antimicrobial treatment leads to increased resistance *in vitro* [79–85], the development of resistance or sensitivity *in vivo* may, in some ways, be a result of stochastic processes or other evolutionary drivers if antibiotic treatment regimens are only exerting weak selective pressure.

It is often assumed that the evolution of AMR involves a fitness cost, although this has predominantly been tested in lab-evolved strains [15, 85–88]. We found no evidence for collateral sensitivity or trade-offs between resistance and fitness in a CF-like medium for these clinical populations. However, in interpreting these results, we must consider that *in vitro* susceptibility and growth testing does not accurately recapitulate the infectious microenvironment of an *in vivo* lung [64]. Therefore, trade-offs between these measures may be present in the lung but not detectable under laboratory conditions. Collateral sensitivity, although shown in evolutionary experiments [15–19], has yet to be demonstrated as widely prevalent in naturally occurring clinical strains. Further work is needed to show that collateral sensitivity is a viable approach for future therapeutic consideration. A recent report found evidence for trade-offs between fitness and multi-drug resistance in clinical *P. aeruginosa* populations [89]. Taken together with our results, we hypothesize that resistance to a single antibiotic may not exert sufficient fitness cost to act as a driving force for trade-offs with growth rate, while resistance to multiple antibiotics perhaps does. Furthermore, this study found stronger evidence for trade-offs in mixed strain infections, whereas all of the individuals in our cohort were infected with a single strain of *P. aeruginosa*. Moreover, as the majority of our strains were technically clinically sensitive to the tested antimicrobials, we may not have had the power to detect trade-offs if they are only elicited at high resistance levels. If resistance does indeed trade-off with fitness, this suggests that slow-growing strains may prove to be the most resistant to treatment. The implication of this for the clinic is concerning, as the slowest growing strains may be more likely to remain undetected during routine susceptibility testing in the clinic, where quick results are favored in order to expedite treatment.

Conducting deep sampling of clinical *P. aeruginosa* populations has allowed us to illuminate population structure, evolution, and population diversity in CF in a manner that single-isolate sampling or population-level sequencing cannot. These methods suffer in their ability to identify rare variants, accurately resolve population structure, and in the case of pooled deep-sequencing, link genotype to phenotype for individual isolates. A 2016 study claimed that single-isolate sampling of longitudinal isolates was sufficient to capture the evolutionary pathways of *P. aeruginosa* in CF lung infection; however, the authors conducted metagenome sequencing at a low depth of 10-31x and only sought to determine if SNPs within individual isolates could be re-discovered in the metagenomes, not whether the individual isolates captured the full diversity of the metagenome [90]. However, we believe there is still incredible value in conducting longitudinal analyses. Building upon previous work [91], we propose that conducting deeper sampling of populations over long time scales will help illuminate the full evolutionary dynamics of *P. aeruginosa* populations in the CF lung and lead to insights that will assist in tackling chronic infections.

## Acknowledgements

For funding, we thank the Georgia Institute of Technology, the Cystic Fibrosis Foundation for grants (DIGGLE18I0 and DIGGLE20G0) to S.P.D. and a fellowship to S.A. (AZIMI18F0), CF@LANTA for a fellowship to S.A. (3206AXB), the National Institutes of Health for a grant (R01AI153116) to S.P.D and T.R., and the National Science Foundation for a fellowship to J.V. (DGE-2039655). Any opinion, findings, and conclusions or recommendations expressed in this material are those of the authors and do not necessarily reflect the views of the funding agencies. Access to the Cystic Fibrosis Biospecimen Registry at Emory Children’s Center for CF and Airways Disease Research was provided through Children’s Healthcare of Atlanta and the Emory University Pediatric CF Discovery Core. We thank Arlene Stecenko and Katy Clemmer for assistance in acquiring sputum samples.

## Author contributions

All authors contributed to research design; J.V. performed research and analyzed data; all authors contributed to writing the paper.

## Supplemental figure legends

**Supplemental Figure 1.**
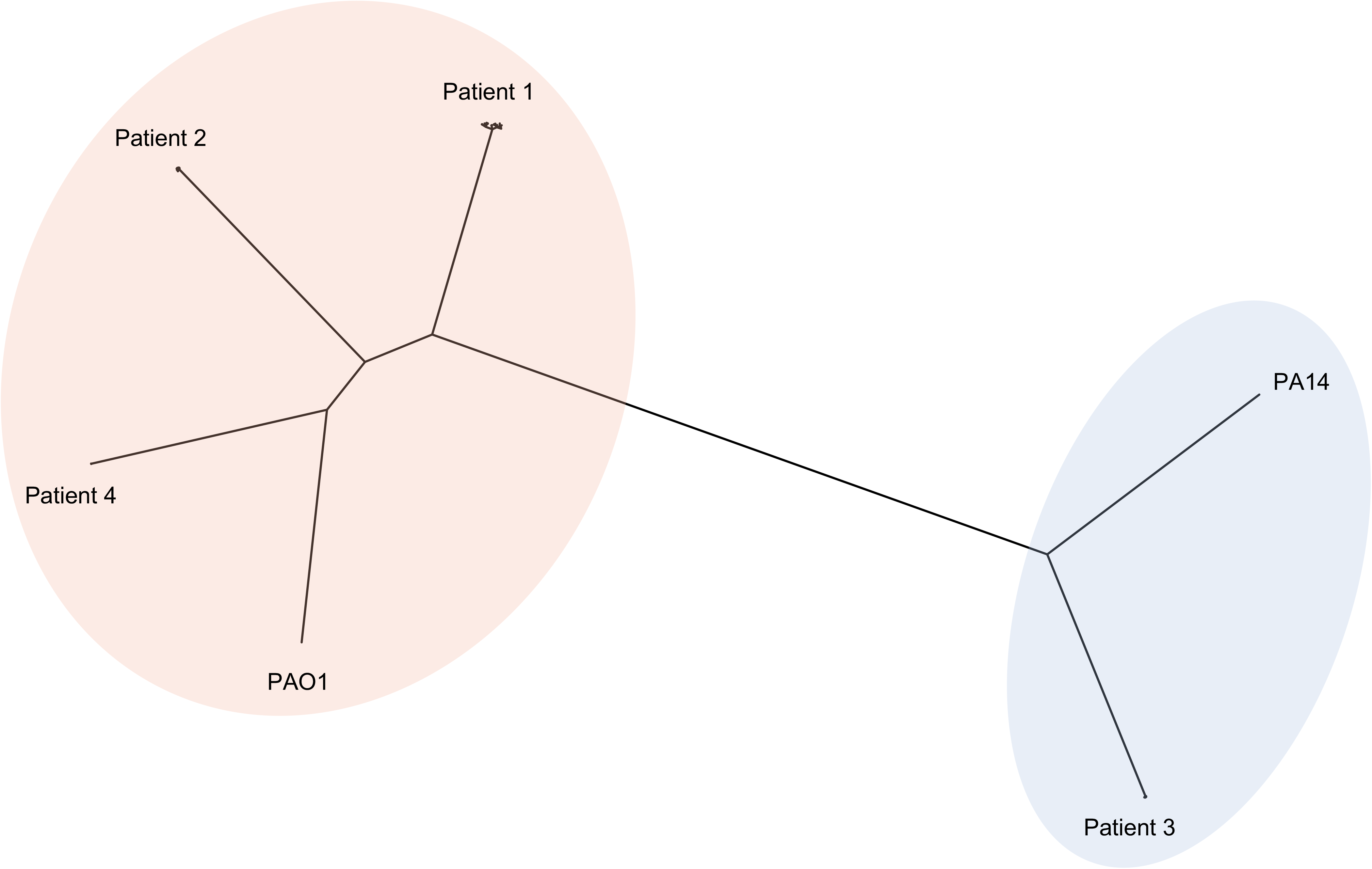
Phylogeny of Patients 1-4 with PAO1 and PA14. Patients 1, 2, and 4 cluster with PAO1, while Patient 3 clusters with PA14.

**Supplemental Figure 2.**
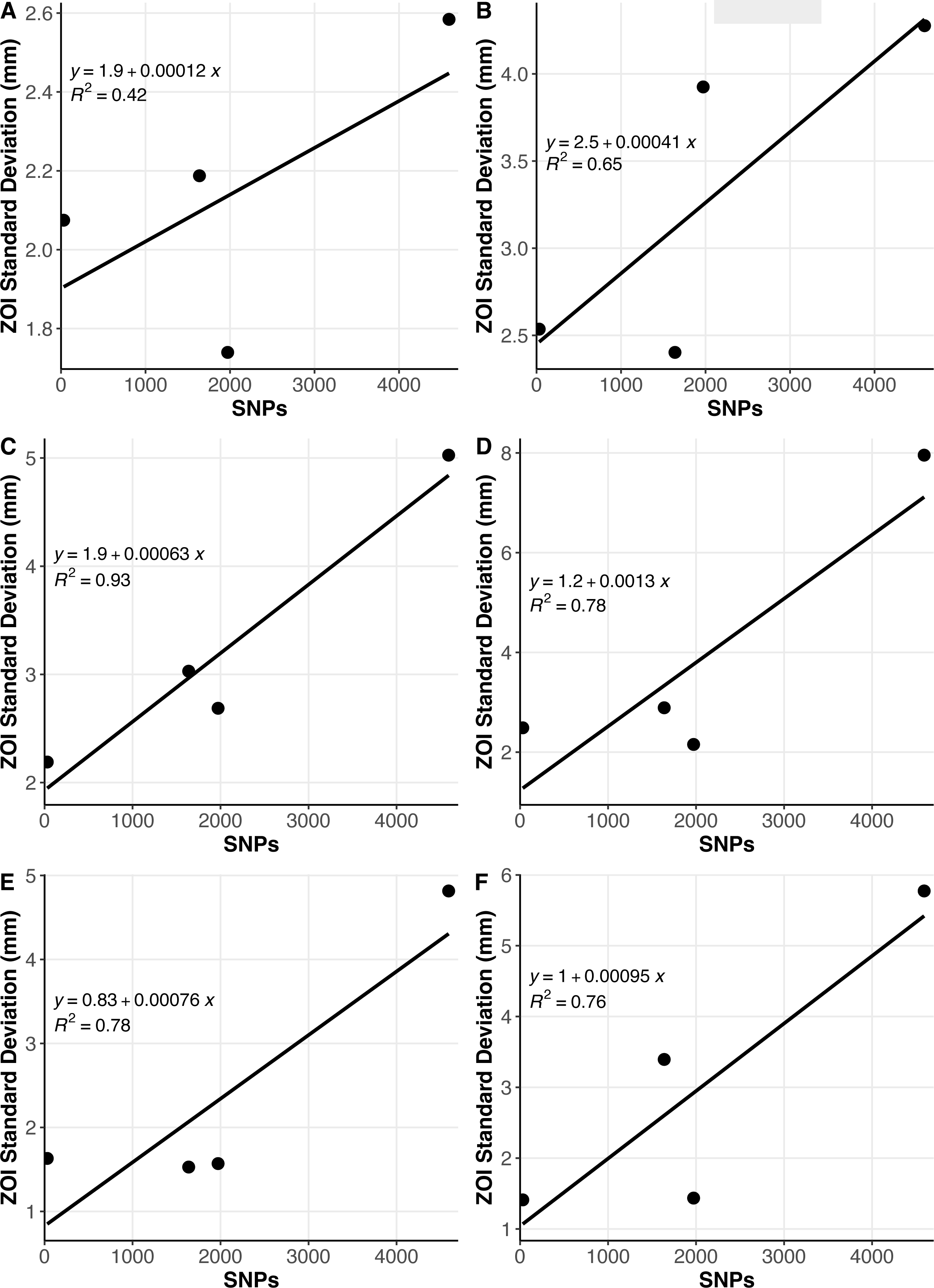
Linear regression analysis demonstrates that total SNP count in a population was a strong indicator of AMR diversity for amikacin (R^2^ = .90, F(1, 2) = 18.94, *p* = .049), meropenem (R^2^ = .93, F(1, 2) = 25.3, *p* = .037), and piperacilin-tazobactam (R^2^ = .95, F(1, 2) = 39.86, *p* = .024), but a poor indicator of AMR diversity for ciprofloxacin (R^2^ = .12, F(1,2) = .27, *p* = .65) and ceftazidime (R^2^ = .71, F(1,2) = 4.78, *p* = .16), and was inversely related to AMR diversity for tobramycin (R^2^ = .97, F(1,2) = 66.61, *p* = .015)

**Supplemental Figure 3.**
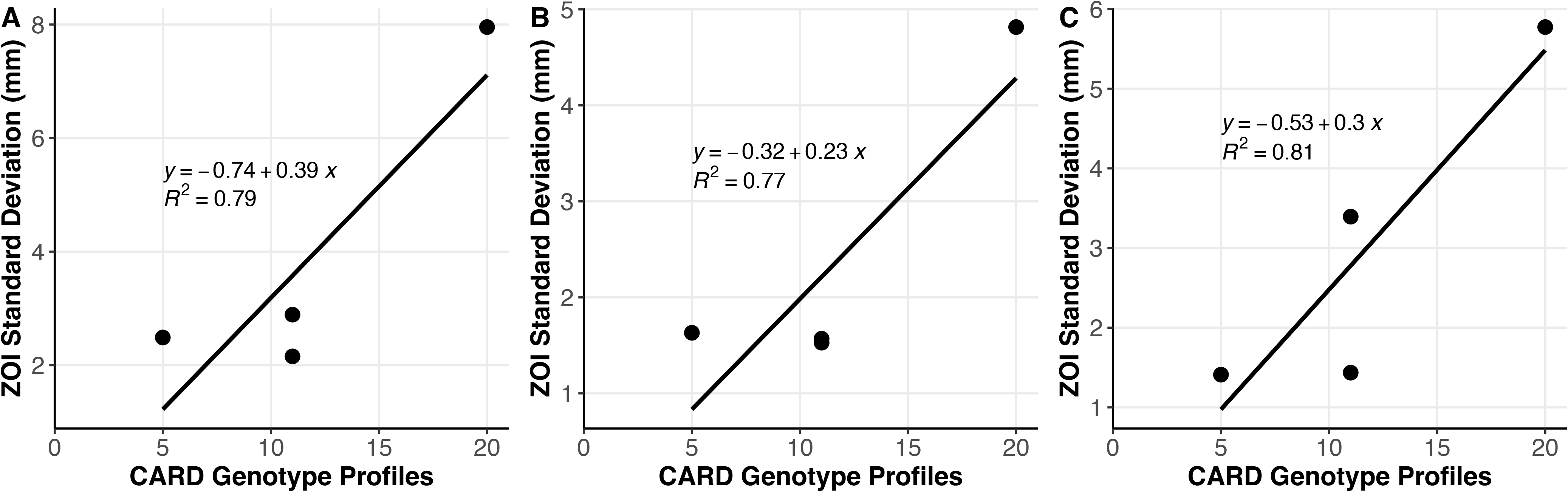
Linear regression analysis shows that the number of distinct CARD profiles within a population is an improved predictor of population standard deviation for ciprofloxacin (R^2^ = .79, F(1,2) = 7.35, *p* = .11), tobramycin (R^2^ = .77, F(1,2) = 6.73, *p* = .12), and ceftazidime (R^2^ = .81, F(1,2) = 8.44, *p* = .10) over total population SNP count.

**Supplemental Figure 4.**
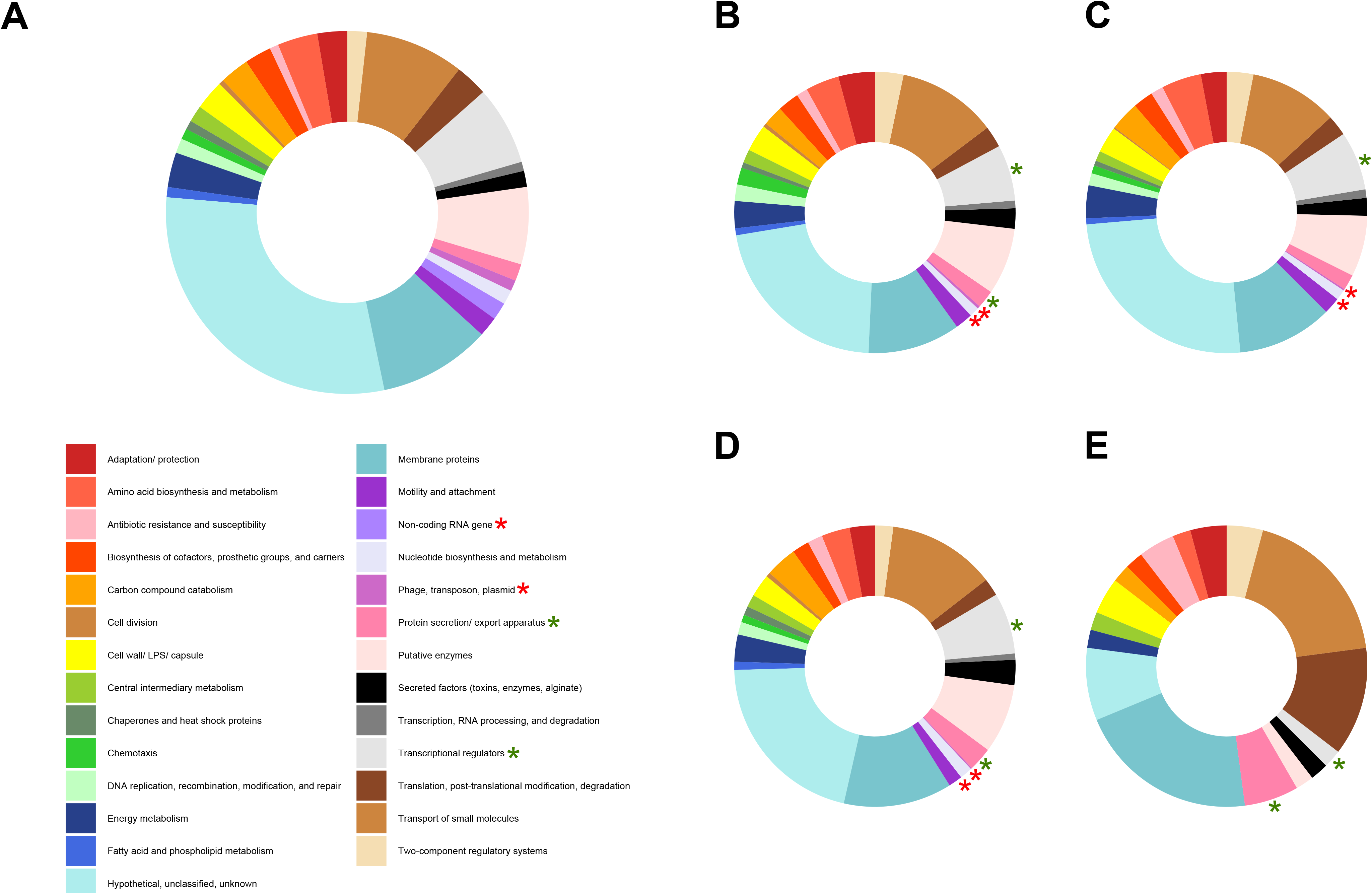
Enrichment analysis of the frequency of functional categories in which non-synonymous SNPs and microindels are found in each of the four populations relative to the proportions of these functional categories in the PAO1 genome shows that protein secretion/ export apparatuses and transcriptional regulators are enriched for such variants, while phage/ transposon/ plasmid and non-coding RNA are less likely to be impacted by such variants. Donut plot of the relative frequencies of genes categorized within each of the 27 different PseudoCAP functional categories in the PAO1 genome (A). Donut plots of the relative frequencies of non-synonymous SNPs and indels located in each of the 27 different PseudoCAP functional categories in Patient 1 (B), 2 (C), 3 (D), and 4 (E). Protein secretion/ export apparatuses and transcriptional regulators are denoted with green asterisks on donut plots where applicable, while phage/ transposon/ plasmid and non-coding RNA are denoted with red asterisks.

**Supplemental Figure 5.**
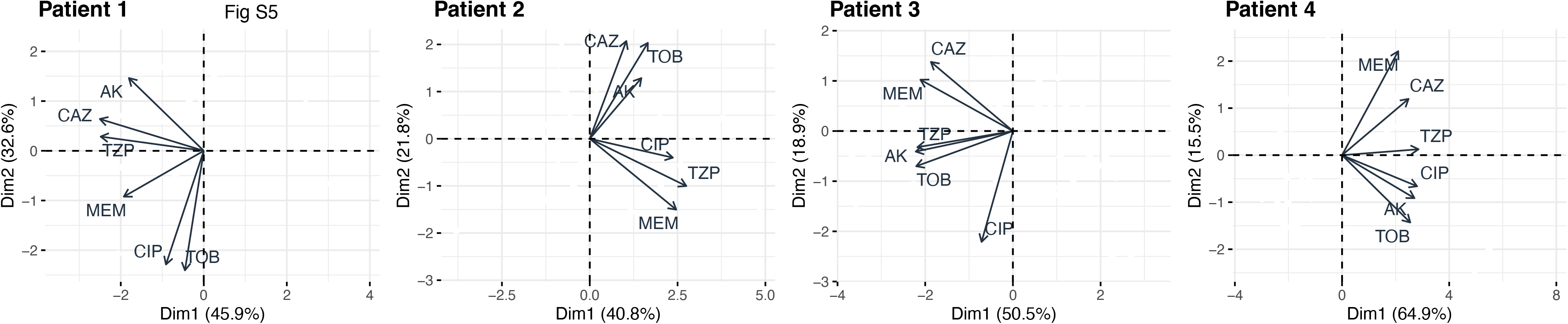
Principal components analysis vectors display no evidence of collateral sensitivity across any of the six antimicrobials tested for any patient, and further demonstrate that cross-resistance and cross-sensitivity patterns differ across patients.

**Supplemental Figure 6.**
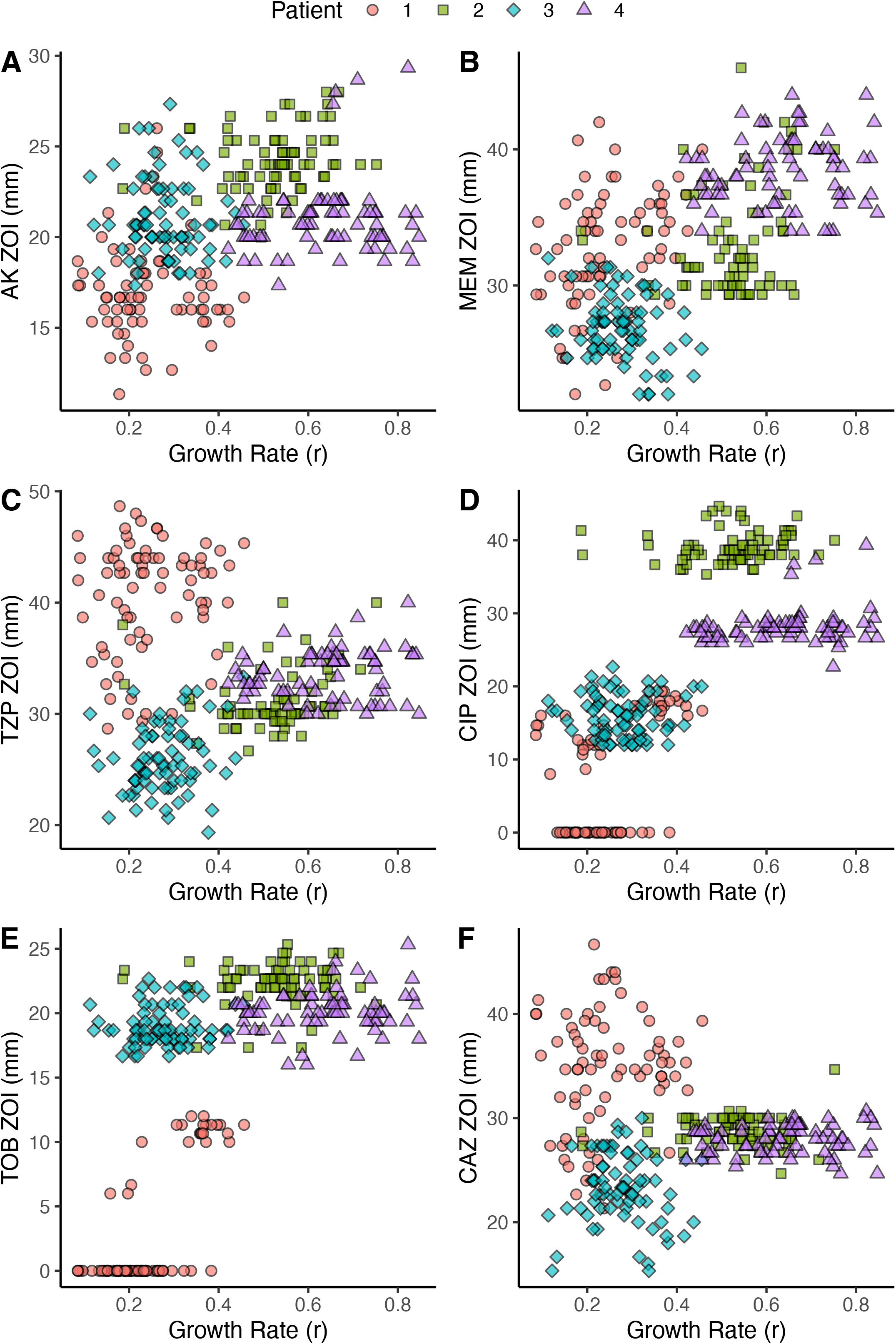
Scatterplots of zone of inhibition (ZOI) versus growth rate (r) in SCFM for all six tested antibiotics: amikacin (AK), meropenem (MEM), piperacillin-tazobactam (TZP), ciprofloxacin (CIP), tobramycin (TOB), and ceftazidime (CAZ). Results of linear mixed model (Table S18), with growth rate in SCFM as a fixed effect and patient as a random effect, demonstrate that there is no significant effect of growth rate on resistance, and therefore, no evidence for trade-offs between growth rate and resistance in these four populations.

**Supplemental Table 1.** Antimicrobial susceptibility testing measurements for Patient 1 as measured by zone of inhibition (ZOI) in a standard disc diffusion assay for amikacin (AK), meropenem (MEM), piperacilin-tazobactam (TZP), ciprofloxacin (CIP), tobramycin (TOB), and ceftazidime (CAZ). Data in the left columns represent raw measurements of zone of inhibition radii (mm units). Data in the right columns represent calculated zone of inhibition values as diameters (mm units).

**Supplemental Table 2.** Antimicrobial susceptibility testing measurements for Patient 2 as measured by zone of inhibition (ZOI) in a standard disc diffusion assay for amikacin (AK), meropenem (MEM), piperacilin-tazobactam (TZP), ciprofloxacin (CIP), tobramycin (TOB), and ceftazidime (CAZ). Data in the left columns represent raw measurements of zone of inhibition radii (mm units). Data in the right columns represent calculated zone of inhibition values as diameters (mm units).

**Supplemental Table 3.** Antimicrobial susceptibility testing measurements for Patient 3 as measured by zone of inhibition (ZOI) in a standard disc diffusion assay for amikacin (AK), meropenem (MEM), piperacilin-tazobactam (TZP), ciprofloxacin (CIP), tobramycin (TOB), and ceftazidime (CAZ). Data in the left columns represent raw measurements of zone of inhibition radii (mm units). Data in the right columns represent calculated zone of inhibition values as diameters (mm units).

**Supplemental Table 4.** Antimicrobial susceptibility testing measurements for Patient 4 as measured by zone of inhibition (ZOI) in a standard disc diffusion assay for amikacin (AK), meropenem (MEM), piperacilin-tazobactam (TZP), ciprofloxacin (CIP), tobramycin (TOB), and ceftazidime (CAZ). Data in the left columns represent raw measurements of zone of inhibition radii (mm units). Data in the right columns represent calculated zone of inhibition values as diameters (mm units).

**Supplemental Table 5.** Genome size, average sequencing coverage, and number of contigs of each assembly.

**Supplemental Table 6.** Supporting statistical values of the linear regression analysis of distinct CARD resistance genotype profiles within a population as a proxy for genomic diversity as measured by total SNPs in each population.

**Supplemental Table 7.** Genes that were impacted by non-synonymous mutations in at least one isolate in all four populations.

**Supplemental Table 8.** Full details of the chi-squared goodness of fit and Monte Carlo simulation exact multinomial tests, with all associated chi-squared and p-values.

**Supplemental Table 9.** Genes that were impacted by non-synonymous mutations in at least one isolate in three out of four populations.

**Supplemental Table 10.** All annotated genetic variants discovered in Patient 1.

**Supplemental Table 11.** All annotated genetic variants discovered in Patient 2.

**Supplemental Table 12.** All annotated genetic variants discovered in Patient 3.

**Supplemental Table 13.** All annotated genetic variants discovered in Patient 4.

**Supplemental Table 14.** Raw OD_600_ reads for growth in SCFM used to create growth curves and analyze growth rate (r) for Patient 1. Time is given in hours, and all isolates were tested in biological triplicates.

**Supplemental Table 15.** Raw OD_600_ reads for growth in SCFM used to create growth curves and analyze growth rate (r) for Patient 2. Time is given in hours, and all isolates were tested in biological triplicates.

**Supplemental Table 16.** Raw OD_600_ reads for growth in SCFM used to create growth curves and analyze growth rate (r) for Patient 3. Time is given in hours, and all isolates were tested in biological triplicates.

**Supplemental Table 17.** Raw OD_600_ reads for growth in SCFM used to create growth curves and analyze growth rate (r) for Patient 4. Time is given in hours, and all isolates were tested in biological triplicates.

**Supplemental Table 18.** Supporting brms R code and statistical values for the linear mixed model run to assess the relationship between growth rate (r) and antimicrobial resistance. Results of the model, with growth rate in SCFM as a fixed effect and patient as a random effect, show that the 95% confidence interval of the fixed effect of growth rate spans 0 for all six antimicrobials. Therefore, the null hypothesis that the fixed effect of growth on antimicrobial susceptibility is 0 cannot be rejected, providing no evidence for trade-offs or any significant relationship between resistance and growth rate across all four populations.

## Notes

### Competing Interest Statement

The authors have declared no competing interest.

### Summary of Updates

Modified the abstract

## References

1. Davies, J.C., *Pseudomonas aeruginosa* in cystic fibrosis: pathogenesis and persistence. Paediatr Respir Rev, 2002. 3(2): p. 128–34.

2. Foundation, C.F., Cystic Fibrosis Foundation Patient Registry 2020 Annual Data Report. 2021.

3. Jackson, L. and V. Waters, Factors influencing the acquisition and eradication of early *Pseudomonas aeruginosa* infection in cystic fibrosis. J Cyst Fibros, 2021. 20(1): p. 8–16.

4. Hoiby, N., O. Ciofu, and T. Bjarnsholt, *Pseudomonas aeruginosa* biofilms in cystic fibrosis. Future Microbiol, 2010. 5(11): p. 1663–74.

5. Van den Bossche, S., et al., The cystic fibrosis lung microenvironment alters antibiotic activity: causes and effects. Eur Respir Rev, 2021. 30(161).

6. Jorth, P., et al., Regional Isolation Drives Bacterial Diversification within Cystic Fibrosis Lungs. Cell Host Microbe, 2015. 18(3): p. 307–19.

7. Clark, S.T., D.S. Guttman, and D.M. Hwang, Diversification of *Pseudomonas aeruginosa* within the cystic fibrosis lung and its effects on antibiotic resistance. FEMS Microbiol Lett, 2018. 365(6).

8. Bartell, J.A., et al., Omics-based tracking of *Pseudomonas aeruginosa* persistence in “eradicated” cystic fibrosis patients. Eur Respir J, 2021. 57(4).

9. Camus, L., F. Vandenesch, and K. Moreau, From genotype to phenotype: adaptations of *Pseudomonas aeruginosa* to the cystic fibrosis environment. Microb Genom, 2021. 7(3).

10. Mehta, H.H., et al., The Essential Role of Hypermutation in Rapid Adaptation to Antibiotic Stress. Antimicrob Agents Chemother, 2019. 63(7).

11. Lujan, A.M., et al., Polymicrobial infections can select against *Pseudomonas aeruginosa* mutators because of quorum-sensing trade-offs. Nat Ecol Evol, 2022. 6(7): p. 979–988.

12. Armbruster, C.R., et al., Heterogeneity in surface sensing suggests a division of labor in *Pseudomonas aeruginosa* populations. Elife, 2019. 8.

13. Porter, S.S. and K.J. Rice, Trade-offs, spatial heterogeneity, and the maintenance of microbial diversity. Evolution, 2013. 67(2): p. 599–608.

14. Ferenci, T., Trade-off Mechanisms Shaping the Diversity of Bacteria. Trends Microbiol, 2016. 24(3): p. 209–223.

15. Barbosa, C., et al., Evolutionary stability of collateral sensitivity to antibiotics in the model pathogen *Pseudomonas aeruginosa*. Elife, 2019. 8.

16. Hernando-Amado, S., F. Sanz-Garcia, and J.L. Martinez, Rapid and robust evolution of collateral sensitivity in *Pseudomonas aeruginosa* antibiotic-resistant mutants. Sci Adv, 2020. 6(32): p. eaba5493.

17. Hernando-Amado, S., et al., Mutational background influences *Pseudomonas aeruginosa* ciprofloxacin resistance evolution but preserves collateral sensitivity robustness. Proc Natl Acad Sci U S A, 2022. 119(15): p. e2109370119.

18. Laborda, P., J.L. Martinez, and S. Hernando-Amado, Convergent phenotypic evolution towards fosfomycin collateral sensitivity of *Pseudomonas aeruginosa* antibiotic-resistant mutants. Microb Biotechnol, 2022. 15(2): p. 613–629.

19. Hernando-Amado, S., et al., Rapid Phenotypic Convergence towards Collateral Sensitivity in Clinical Isolates of *Pseudomonas aeruginosa* Presenting Different Genomic Backgrounds. Microbiol Spectr, 2023. 11(1): p. e0227622.

20. Cramer, N., et al., Microevolution of the major common *Pseudomonas aeruginosa* clones C and PA14 in cystic fibrosis lungs. Environ Microbiol, 2011. 13(7): p. 1690–704.

21. Marvig, R.L., et al., Genome analysis of a transmissible lineage of pseudomonas aeruginosa reveals pathoadaptive mutations and distinct evolutionary paths of hypermutators. PLoS Genet, 2013. 9(9): p. e1003741.

22. Feliziani, S., et al., Coexistence and within-host evolution of diversified lineages of hypermutable *Pseudomonas aeruginosa* in long-term cystic fibrosis infections. PLoS Genet, 2014. 10(10): p. e1004651.

23. Markussen, T., et al., Environmental heterogeneity drives within-host diversification and evolution of *Pseudomonas aeruginosa*. mBio, 2014. 5(5): p. e01592–14.

24. Marvig, R.L., et al., Within-host microevolution of *Pseudomonas aeruginosa* in Italian cystic fibrosis patients. BMC Microbiol, 2015. 15: p. 218.

25. Marvig, R.L., et al., Convergent evolution and adaptation of *Pseudomonas aeruginosa* within patients with cystic fibrosis. Nat Genet, 2015. 47(1): p. 57–64.

26. Bianconi, I., et al., Persistence and Microevolution of *Pseudomonas aeruginosa* in the Cystic Fibrosis Lung: A Single-Patient Longitudinal Genomic Study. Front Microbiol, 2018. 9: p. 3242.

27. Klockgether, J., et al., Long-Term Microevolution of *Pseudomonas aeruginosa* Differs between Mildly and Severely Affected Cystic Fibrosis Lungs. Am J Respir Cell Mol Biol, 2018. 59(2): p. 246–256.

28. Gabrielaite, M., et al., Gene Loss and Acquisition in Lineages of *Pseudomonas aeruginosa* Evolving in Cystic Fibrosis Patient Airways. mBio, 2020. 11(5).

29. Datar, R., et al., Phenotypic and Genomic Variability of Serial Peri-Lung Transplantation *Pseudomonas aeruginosa* Isolates From Cystic Fibrosis Patients. Front Microbiol, 2021. 12: p. 604555.

30. Wardell, S.J.T., et al., Genome evolution drives transcriptomic and phenotypic adaptation in *Pseudomonas aeruginosa* during 20 years of infection. Microb Genom, 2021. 7(11).

31. Andersson, D.I., H. Nicoloff, and K. Hjort, Mechanisms and clinical relevance of bacterial heteroresistance. Nat Rev Microbiol, 2019. 17(8): p. 479–496.

32. Reece, E., P.H.A. Bettio, and J. Renwick, Polymicrobial Interactions in the Cystic Fibrosis Airway Microbiome Impact the Antimicrobial Susceptibility of *Pseudomonas aeruginosa*. Antibiotics (Basel), 2021. 10(7).

33. Galdino, A.C.M., et al., Polymicrobial Biofilms in Cystic Fibrosis Lung Infections: Effects on Antimicrobial Susceptibility, in Multispecies Biofilms: Technologically Advanced Methods to Study Microbial Communities, K.S. Kaushik and S.E. Darch, Editors. 2023, Springer International Publishing: Cham. p. 231–267.

34. Mowat, E., et al., *Pseudomonas aeruginosa* population diversity and turnover in cystic fibrosis chronic infections. Am J Respir Crit Care Med, 2011. 183(12): p. 1674–9.

35. Ashish, A., et al., Extensive diversification is a common feature of *Pseudomonas aeruginosa* populations during respiratory infections in cystic fibrosis. J Cyst Fibros, 2013. 12(6): p. 790–3.

36. Workentine, M.L., et al., Phenotypic heterogeneity of *Pseudomonas aeruginosa* populations in a cystic fibrosis patient. PLoS One, 2013. 8(4): p. e60225.

37. Clark, S.T., et al., Phenotypic diversity within a *Pseudomonas aeruginosa* population infecting an adult with cystic fibrosis. Sci Rep, 2015. 5: p. 10932.

38. Darch, S.E., et al., Recombination is a key driver of genomic and phenotypic diversity in a *Pseudomonas aeruginosa* population during cystic fibrosis infection. Sci Rep, 2015. 5: p. 7649.

39. Williams, D., et al., Divergent, coexisting *Pseudomonas aeruginosa* lineages in chronic cystic fibrosis lung infections. Am J Respir Crit Care Med, 2015. 191(7): p. 775–85.

40. Williams, D., et al., Transmission and lineage displacement drive rapid population genomic flux in cystic fibrosis airway infections of a *Pseudomonas aeruginosa* epidemic strain. Microb Genom, 2018. 4(3).

41. Su, M., S.W. Satola, and T.D. Read, Genome-Based Prediction of Bacterial Antibiotic Resistance. J Clin Microbiol, 2019. 57(3).

42. Palmer, K.L., L.M. Aye, and M. Whiteley, Nutritional cues control *Pseudomonas aeruginosa* multicellular behavior in cystic fibrosis sputum. J Bacteriol, 2007. 189(22): p. 8079–87.

43. Petit, R.A., 3rd and T.D. Read, Bactopia: a Flexible Pipeline for Complete Analysis of Bacterial Genomes. mSystems, 2020. 5(4).

44. Jolley, K.A., J.E. Bray, and M.C.J. Maiden, Open-access bacterial population genomics: BIGSdb software, the PubMLST.org website and their applications. Wellcome Open Res, 2018. 3: p. 124.

45. Wick, R.R., et al., Unicycler: Resolving bacterial genome assemblies from short and long sequencing reads. PLoS Comput Biol, 2017. 13(6): p. e1005595.

46. Research, O.N.T., Medaka. 2018, Oxford Nanopore Technologies.

47. Wick, R.R. and K.E. Holt, Polypolish: Short-read polishing of long-read bacterial genome assemblies. PLoS Comput Biol, 2022. 18(1): p. e1009802.

48. Walker, B.J., et al., Pilon: an integrated tool for comprehensive microbial variant detection and genome assembly improvement. PLoS One, 2014. 9(11): p. e112963.

49. Seemann, T., Snippy. 2020.

50. Seemann, T., Prokka: rapid prokaryotic genome annotation. Bioinformatics, 2014. 30(14): p. 2068–9.

51. Otto, T.D., et al., RATT: Rapid Annotation Transfer Tool. Nucleic Acids Res, 2011. 39(9): p. e57.

52. Oliver, A., et al., High frequency of hypermutable *Pseudomonas aeruginosa* in cystic fibrosis lung infection. Science, 2000. 288(5469): p. 1251–4.

53. Alcock, B.P., et al., CARD 2023: expanded curation, support for machine learning, and resistome prediction at the Comprehensive Antibiotic Resistance Database. Nucleic Acids Res, 2023. 51(D1): p. D690–D699.

54. Engels, B., XNomial. 2015.

55. Croucher, N.J., et al., Rapid phylogenetic analysis of large samples of recombinant bacterial whole genome sequences using Gubbins. Nucleic Acids Res, 2015. 43(3): p. e15.

56. Sprouffske, K. and A. Wagner, Growthcurver: an R package for obtaining interpretable metrics from microbial growth curves. BMC Bioinformatics, 2016. 17: p. 172.

57. Bürkner, P., brms: An R Package for Bayesian Multilevel Models using Stan. Journal of Statistical Software, 2017. 80(1): p. 1–28.

58. Toledo-Arana, A., F. Repoila, and P. Cossart, Small noncoding RNAs controlling pathogenesis. Curr Opin Microbiol, 2007. 10(2): p. 182–8.

59. Hernando-Amado, S., et al., Rapid Decline of Ceftazidime Resistance in Antibiotic-Free and Sublethal Environments Is Contingent on Genetic Background. Mol Biol Evol, 2022. 39(3).

60. Genova, R., et al., Collateral Sensitivity to Fosfomycin of Tobramycin-Resistant Mutants of Pseudomonas aeruginosa Is Contingent on Bacterial Genomic Background. Int J Mol Sci, 2023. 24(8).

61. Roemhild, R. and D.I. Andersson, Mechanisms and therapeutic potential of collateral sensitivity to antibiotics. PLoS Pathog, 2021. 17(1): p. e1009172.

62. Jean-Pierre, F., et al., Community composition shapes microbial-specific phenotypes in a cystic fibrosis polymicrobial model system. Elife, 2023. 12.

63. Somayaji, R., et al., Antimicrobial susceptibility testing (AST) and associated clinical outcomes in individuals with cystic fibrosis: A systematic review. J Cyst Fibros, 2019. 18(2): p. 236–243.

64. Bjarnsholt, T., et al., The importance of understanding the infectious microenvironment. Lancet Infect Dis, 2022. 22(3): p. e88–e92.

65. Macia, M.D., et al., Hypermutation is a key factor in development of multiple-antimicrobial resistance in *Pseudomonas aeruginosa* strains causing chronic lung infections. Antimicrob Agents Chemother, 2005. 49(8): p. 3382–6.

66. Cabot, G., et al., Evolution of *Pseudomonas aeruginosa* Antimicrobial Resistance and Fitness under Low and High Mutation Rates. Antimicrob Agents Chemother, 2016. 60(3): p. 1767–78.

67. Khil, P.P., et al., Dynamic Emergence of Mismatch Repair Deficiency Facilitates Rapid Evolution of Ceftazidime-Avibactam Resistance in *Pseudomonas aeruginosa* Acute Infection. mBio, 2019. 10(5).

68. Rees, V.E., et al., Characterization of Hypermutator *Pseudomonas aeruginosa* Isolates from Patients with Cystic Fibrosis in Australia. Antimicrob Agents Chemother, 2019. 63(4).

69. Colque, C.A., et al., Hypermutator Pseudomonas aeruginosa Exploits Multiple Genetic Pathways To Develop Multidrug Resistance during Long-Term Infections in the Airways of Cystic Fibrosis Patients. Antimicrob Agents Chemother, 2020. 64(5).

70. Colque, C.A., et al., Longitudinal Evolution of the Pseudomonas-Derived Cephalosporinase (PDC) Structure and Activity in a Cystic Fibrosis Patient Treated with beta-Lactams. mBio, 2022. 13(5): p. e0166322.

71. Dulanto Chiang, A., et al., Hypermutator strains of *Pseudomonas aeruginosa* reveal novel pathways of resistance to combinations of cephalosporin antibiotics and beta-lactamase inhibitors. PLoS Biol, 2022. 20(11): p. e3001878.

72. Moriarty, T.F., et al., Sputum antibiotic concentrations: implications for treatment of cystic fibrosis lung infection. Pediatr Pulmonol, 2007. 42(11): p. 1008–17.

73. Ciofu, O., et al., Antimicrobial resistance, respiratory tract infections and role of biofilms in lung infections in cystic fibrosis patients. Adv Drug Deliv Rev, 2015. 85: p. 7–23.

74. Sonderholm, M., et al., The Consequences of Being in an Infectious Biofilm: Microenvironmental Conditions Governing Antibiotic Tolerance. Int J Mol Sci, 2017. 18(12).

75. Crabbe, A., et al., Antimicrobial Tolerance and Metabolic Adaptations in Microbial Biofilms. Trends Microbiol, 2019. 27(10): p. 850–863.

76. Martin, I., V. Waters, and H. Grasemann, Approaches to Targeting Bacterial Biofilms in Cystic Fibrosis Airways. Int J Mol Sci, 2021. 22(4).

77. Santi, I., et al., Evolution of Antibiotic Tolerance Shapes Resistance Development in Chronic Pseudomonas aeruginosa Infections. mBio, 2021. 12(1).

78. Witzany, C., R.R. Regoes, and C. Igler, Assessing the relative importance of bacterial resistance, persistence and hyper-mutation for antibiotic treatment failure. Proc Biol Sci, 2022. 289(1986): p. 20221300.

79. Wong, A., N. Rodrigue, and R. Kassen, Genomics of adaptation during experimental evolution of the opportunistic pathogen *Pseudomonas aeruginosa*. PLoS Genet, 2012. 8(9): p. e1002928.

80. Jorgensen, K.M., et al., Sublethal ciprofloxacin treatment leads to rapid development of high-level ciprofloxacin resistance during long-term experimental evolution of *Pseudomonas aeruginosa*. Antimicrob Agents Chemother, 2013. 57(9): p. 4215–21.

81. Jorth, P., et al., Evolved Aztreonam Resistance Is Multifactorial and Can Produce Hypervirulence in *Pseudomonas aeruginosa*. mBio, 2017. 8(5).

82. Ahmed, M.N., et al., Evolution of Antibiotic Resistance in Biofilm and Planktonic *Pseudomonas aeruginosa* Populations Exposed to Subinhibitory Levels of Ciprofloxacin. Antimicrob Agents Chemother, 2018. 62(8).

83. Sanz-Garcia, F., S. Hernando-Amado, and J.L. Martinez, Mutation-Driven Evolution of *Pseudomonas aeruginosa* in the Presence of either Ceftazidime or Ceftazidime-Avibactam. Antimicrob Agents Chemother, 2018. 62(10).

84. Wardell, S.J.T., et al., A large-scale whole-genome comparison shows that experimental evolution in response to antibiotics predicts changes in naturally evolved clinical *Pseudomonas aeruginosa*. Antimicrob Agents Chemother, 2019. 63(12).

85. Ahmed, M.N., et al., The evolutionary trajectories of *Pseudomonas aeruginosa* in biofilm and planktonic growth modes exposed to ciprofloxacin: beyond selection of antibiotic resistance. NPJ Biofilms Microbiomes, 2020. 6(1): p. 28.

86. Hernando-Amado, S., et al., Fitness costs associated with the acquisition of antibiotic resistance. Essays Biochem, 2017. 61(1): p. 37–48.

87. Laborda, P., J.L. Martinez, and S. Hernando-Amado, Evolution of Habitat-Dependent Antibiotic Resistance in *Pseudomonas aeruginosa*. Microbiol Spectr, 2022. 10(4): p. e0024722.

88. Jorth, P., et al., Evolved bacterial siderophore-mediated antibiotic cross-protection. Preprint available at 10.21203/rs.3.rs-2644953/v1, 2023.

89. Diaz Caballero, J., et al., Mixed strain pathogen populations accelerate the evolution of antibiotic resistance in patients. Nat Commun, 2023. 14(1): p. 4083.

90. Sommer, L.M., et al., Is genotyping of single isolates sufficient for population structure analysis of *Pseudomonas aeruginosa* in cystic fibrosis airways? BMC Genomics, 2016. 17: p. 589.

91. Diaz Caballero, J., et al., Selective Sweeps and Parallel Pathoadaptation Drive *Pseudomonas aeruginosa* Evolution in the Cystic Fibrosis Lung. mBio, 2015. 6(5): p. e00981–15.

